# Long range segmentation of prokaryotic genomes by gene age and functionality

**DOI:** 10.1101/2024.04.26.591304

**Authors:** Yuri I. Wolf, Ilya V. Schurov, Kira S. Makarova, Mikhail I. Katsnelson, Eugene V. Koonin

## Abstract

Bacterial and archaeal genomes encompass numerous operons that typically consist of two to five genes. On larger scales, however, gene order is poorly conserved through the evolution of prokaryotes. Nevertheless, non-random localization of different classes of genes on prokaryotic chromosomes could reflect important functional and evolutionary constraints. We explored the patterns of genomic localization of evolutionarily conserved (ancient) and variable (young) genes across the diversity of bacteria and archaea. Nearly all bacterial and archaeal chromosomes were found to encompass large segments of 100-300 kilobases that were significantly enriched in either ancient or young genes. Similar clustering of genes with lethal knockout phenotype (essential genes) was observed as well. Mathematical modeling of genome evolution suggests that this long-range gene clustering in prokaryotic chromosomes reflects perpetual genome rearrangement driven by a combination of selective and neutral processes rather than evolutionary conservation.

## Introduction

Genome evolution in prokaryotes (bacteria and archaea) is a highly dynamic process that involves extensive gene gain via horizontal gene transfer (HGT), gene loss, and genome rearrangement via inversion and translocation of genome segments (1–3). Prokaryotic pangenomes, that is, the entirety of the genes in a species, consist of the core genome, i.e. the set of conserved genes that are shared by all strains in a species, and the accessory genome consisting of genes that are rapidly replaced and exchanged during evolution (4–6). Functionally associated genes in prokaryotes typically form operons, that is, co-transcribed arrays that typically consist of 2 to 5 genes, with the exception of several longer ones, such as the ribosomal super-operon (7–9). Operons are often shared by genomes of distantly related prokaryotes, partly due to evolutionary conservation maintained by selective pressure for coregulation of functionally linked genes and partly to HGT, under the selfish operon hypothesis (10,11).

In addition to operonic organization, it has been noticed that genes comprising the core genome (persistent genes) tend to cluster in bacterial genomes, and the clusters of these genes only partially corresponded to operons (12). This clustering has been attributed to selection for minimization of the disruption of functionally important genes by deletions.

Beyond the operon scale, gene order in prokaryotic genomes is poorly conserved such that genomic synteny is lost even in closely related bacteria and archaea that belong to the same genus and share high sequence similarity between orthologous genes (13,14). Genomic synteny is disrupted by multiple processes including insertion of DNA via HGT, deletions and frequent inversion of genome segments that tends to be symmetrical with respect to the origin of replication (1,15).

Notwithstanding the weak conservation of genomic synteny, there are various indications of non-randomness in long range gene localization in genomes of prokaryotes (16). A prominent feature of many bacterial genomes is the presence of various genomic islands, such as defense islands enriched in antivirus defense systems (17) and pathogenicity islands carrying various virulence genes (18–20). Genomic islands vary in size from several to hundreds of genes and are both enriched in various integrated mobile genetic elements including plasmids, integrative conjugating elements and viruses, and mobilized by these elements (21,22).

A different type of regularity is observed in gene organization in bacterial chromosomes with respect to the origin of DNA replication whereby highly expressed and essential genes are located primarily on the leading DNA strand (23–25). This strand bias in gene localization is generally thought to be caused by selection against deleterious collisions between the replisome and the transcribing RNA polymerase although alternative explanations of this phenomenon have been proposed as well.

Non-random gene organization has been associated also with three-dimensional structure of prokaryotic chromosomes. In particular, it has been found that spatial domains in bacterial chromosomes (chromosomal interaction domains) are bounded by long, highly expressed genes (26,27). Most conspicuous is the case of the archaea of the genus *Sulfolobus*, in which the chromosome is partitioned into two distinct spatial compartments, A and B, with the A compartment being notably enriched in highly expressed, evolutionarily conserved genes (28).

We explored the patterns of genomic localization of evolutionarily conserved (ancient) and variable (young) genes across the diversity of bacteria and archaea. In nearly all analyzed bacterial and archaeal chromosomes, we identified large segments of 100-300 kilobases (kb) that were significantly enriched in either ancient or young genes. Similar clustering of genes with lethal knockout phenotype (essential genes) was observed as well. This mesoscale gene clustering in prokaryotic chromosomes appears to reflect perpetual genome rearrangement driven by a combination of selective and neutral processes rather than evolutionary conservation.

## Materials and Methods

### Genome sets of prokaryote genera

A collection of completely sequenced bacterial and archaeal genomes was downloaded from NCBI Genomes (https://ftp.ncbi.nlm.nih.gov/genomes/ASSEMBLY_REPORTS/) in November 2021 (29). According to the NCBI taxonomy (as of November 2021), it included 175 genera that were each represented by at least 15 assemblies.

For each of the 175 microbial genera, the entire complement of (predicted) protein sequences was collected and clustered using MMSEQS2 (30) with the similarity threshold of 0.5. The resulting protein clusters were mapped back to the assemblies, such that each assembly could be represented by a set of cluster IDs.

First, redundant assemblies (identical with respect to 0.5 similarity clusters), were identified and discarded. Second, for each protein cluster, the number of assemblies that harbored at least one representative protein was determined. The distribution of these numbers (the gene commonality curve) typically has an asymmetrical U-shape (1), and this property was used to detect outliers in ostensibly genus-wide genome sets. To this end, the rightmost peak of the commonality plot was identified, and only those assemblies were that contained at least one representative of each of the clusters in the peak bin. Formally, for a *k*-assembly set with the *n_i_* denoting the number of clusters that are present in exactly *i* genomes, 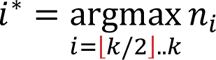 was found. Then, the list of clusters, present in exactly *i*^∗^ assemblies, was used to screen the full set of *k* assemblies, reducing it to *i*^∗^ assemblies. Then, the commonality curve was recalculated for the 0.5 similarity clusters and the new assembly set.

This procedure yielded a total of 20,207 genome assemblies representing 175 genera, with 12 to 1,879 assemblies per genus.

### Operational classification of genes

The commonality distribution of 0.5 similarity clusters was used to classify all genes within each genus into three classes: least common, most common, and intermediate. This classification requires two commonality thresholds to separate the classes. To obtain these thresholds, the commonality list for all clusters encoded in each genome was compiled and the inclusive quantiles for the top and bottom 1/3rds of the distribution were determined (S1 Figure, S8 Figure). This procedure was performed for the entire genus-wide set of assemblies, and the quantiles 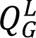 and 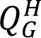 were used as the thresholds for the given genus, and for each genome separately; genomes with individual quantiles 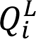 and 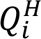 were ranked according to its similarity to the genus-vide value, 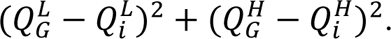

For each genome, genes with the commonality of the corresponding 0.5 similarity cluster greater than or equal to 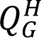 were classified into the “most common” category, and genes with the commonality below 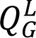 were classified as the “least common”. These gene classes were operationally defined as “ancient” and “young” genes, respectively.

### Different levels of evolutionary conservation

For two model organisms, the archaeon *Sulfolobus islandicus* M.16.4 (*Thermoproteota, Sulfolobales*) and the bacterium *Escherichia coli* K-12 sub. BW25113 (*Pseudomonadota, Enterobacterales*) we considered additional levels of evolutionary conservation. The proteins encoded in these two genomes were mapped to arCOGs and COGs, respectively (31,32). For *S. islandicus*, arCOGs with the posterior probability of presence in the last archaeal common ancestor (33) of at least 0.95 were considered to represent the deep level of evolutionary conservation and arCOGs present in at least 75% of the genomes of *Desulfurococcales*, *Acidilobales*, *Fervidicoccales*, and *Sulfolobales* were considered to represent the intermediate level of evolutionary conservation. For *E. coli*, COGs present in at least 15 of the 20 bacterial phyla were considered to represent the deep level of evolutionary conservation and COGs present in at least 75% of the genomes of *Pseudomonadota* (formerly, Proteobacteria) were considered to represent the intermediate level of evolutionary conservation.

### Segmentation of genome partitions

A genome partition, such as a chromosome, can be represented as a string of genes. Any binary attribute assigned to a gene (e.g. whether the corresponding 0.5 similarity protein cluster is “young” or whether the knockout of this gene has the lethal phenotypic effect) thus defines a binary sequence representing the mapping of this attribute to the chromosome. To account for the circularity of the major genome partitions in prokaryotes, the genome midpoint *K* = ⌊*N*/2⌋ where *N* is the number of genes in the partition was identified and the genome was duplicated, to form the sequence of domain attributes *a_i_* with indices in the range (1. .2*N*), corresponding to the original indices as ((*K* + 1). . *N*, 1. . *N*, 1. . *K*). Then, a skew profile was calculated as *s_i_* = *s_i-1_* + *a_i_* − *M*/*N* (formally, setting *s*_0_ =0), where 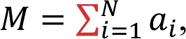 i.e. the sum of attributes across the genome. Rising segments of this profile indicate regions where the local density of the attribute is greater than the genome average, whereas falling segments indicate below the average regions. Gaussian kernel smoothing with a bandwidth *b* = 40 domains (formally, 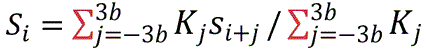 where *K_j_* = exp (−(*j*/*b*)^2^/2) is the observation weight at the distance *j* from *i*-th position) was applied to the skew profile to eliminate small-scale variations. Minima and maxima of the smoothed profile *S_i_*, corresponding to the original partition (indexes ((*N* − *K* + 1). . (2*N* − *K*)) in the duplicated partition) provide the initial segmentation of the genome into alternating regions with above- and below-average densities of the given attribute.

This segmentation was further refined using the following greedy procedure. First, consider a current segmentation of the chromosome into *k* segments where the *i*-th segment contains *n_i_* genes, *m_i_* of which have the attribute value of 1 and *n_i_* − *m_i_* have the attribute value of 0, so that the attribute density in this segment is *р_i_* = *m_i_*/*n_i_*. The log likelihood of this segment can be calculated from the binomial probability as *L_i_* = *m_i_* ln *р_i_* + (*n_i_* − *m_i_*) ln(1 − *р_i_*) and the global log likelihood as 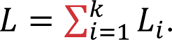 Such a segmentation of a circular genome partition depends on *k* segment boundaries and *k* − 1 independent segment densities (the global average attribute density serves as a constraint). The global measure of the “information cost” of such segmentation is given by the Bayesian Information Criterion, *B* = (2*k* − 1) ln *N* − 2*L*.

Now, consider merging segment *i* + 1 into segment *i*. The merged segment will have 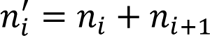 domains, 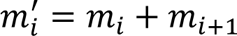 of which are positive for the attribute. The log likelihood for the merged segment 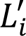 can be calculated from 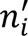 and 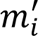 as above, whereas 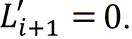 This results in the total log likelihood of *L*^i^ (denoting the likelihood after the elimination of segment *i* + 1) and the “information cost” of *B*^i^ = (2*k* − 3) ln *N* − 2*L*^i^ (elimination of one segment reduces the number of parameters by two). The best available merging is defined by 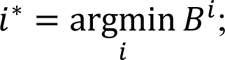 if *B*^i∗^ < *B* (improving the total “information cost” relative to the original segmentation), the segments *i*^∗^and *i*^∗^ + 1 are merged and the procedure is repeated until convergence.

The robustness of the approach to random fluctuations was tested using artificial chromosomes with permuted gene order; the algorithm fails to segment such randomized gene strings and reports a single segment with uniform density.

### Statistical test of genome segmentation

Consider a chromosome consisting of *N* genes, of which *M* are tagged (i.e. labeled “young”, “ancient”, or with any other attribute). Consider a segmentation of the chromosome into *k* segments where the *i*-th segment contains *n_i_* genes, *m_i_*of which are tagged (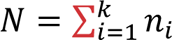 and 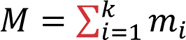). If the density of tagged genes is uniform, the expected number of tagged genes in each segment can be calculated as 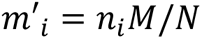 and the deviation of the observed distribution from the expectation is characterized by the 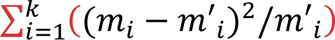 statistics, which follows the *χ*^2^ distribution with *k* − 1 degrees of freedom.

### Multiscale structural complexity

To visualize the structure of a tagged chromosome, a multiscale structural complexity (34) profile was constructed as follows. The chromosome is modeled as a circular sequence (*x*_1_, *x*_2_, …, *x_n_*) of 0’s and 1’s, where 0 corresponds to a gene lacking a given attribute and 1 corresponds to a gene possessing that attribute. For each scale level *λ* from 1 to the half of the number of genes in the chromosome, a smoothed version of the chromosome was obtained by calculating a rolling mean with a window of length *λ*, that is, a new circular sequence (*y*_1_, *y*_2_, …, *y_n_*) where 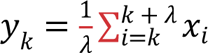 was obtained. Then, for each *λ*, the average absolute difference between the chromosomes smoothed at levels *λ* and 2*λ* was calculated. The plot of this difference as a function of *λ* is the multiscale structural complexity profile.

The idea behind this construction is to characterize the contribution of features of a certain characteristic size to the overall variance of the pattern. For example, if 0’s and 1’s are distributed randomly (by coin-flipping), the pattern will be smoothed out to -near uniformity even for relatively small values of *λ*, and thus there would be little difference between smoothing at levels *λ* and 2*λ*, except for very small values of *λ*. Therefore, in the complexity profile, there will be a sharp peak at small values of *λ*. By contrast, if the pattern consists of large contiguous blocks of 0’s and 1’s, smoothing results at levels *λ* and 2*λ* will substantially differ for small values of *λ*, and the maximum of the complexity profile will lie near the characteristic size of the blocks. If there is a combination of the two types of patterns, that is, large blocks distorted by noise, the profile will be multimodal.

In this work, multiscale complexity profiles were used to qualitatively and quantitatively compare tagged chromosomes, both simulated and real. Specifically, the mean absolute difference between the complexity profiles was used as a measure of the distance between chromosomes.

### Fitting of the differential mobility model

To fit the parameters of the model evolution to a given real genome, a combination of random and grid search was performed as follows. The chromosome size and the number of mobile genes are taken from the real genome, and the width of the attractor basin *γ* is fixed at 1. The parameters being optimized are the number of attractors *a*, the mobility bonus *β* and the attractor strength. The number of attractors *a* is an integer changing in the range [1, 30]. To explore the ranges of values of *β* and *α* over several orders of magnitude, the following parametrization was used: *β* =*exp β*′ and *α* =*exp* (*α*′) − 1, and with *β*′ spanning the range [*log* 0.001,*log* 30] and *α*′ spanning the range [*log* 0.1,*log* 1000].

At the first stage, a random search was performed by sampling 2000 values of (*a*, *β*′, *α*′) taken from the uniform distribution over the ranges indicated above. For each value of the parameters, the simulation started from a random genome, and then, 500 epochs of virtual evolution were run, each epoch consisting of 5,000 evolution steps. At the end of each epoch, the binary sequence, representing low- and high-mobility genes in the simulated genome, was recorded.

Then, the mean absolute difference between the complexity profiles of each of the recorded simulated genomes and the real genome was calculated, and the bottom 0.1-quantile of these distances was recorded as a goodness-of-fit measure (smaller values indicate better fit) value for a given combination of the parameter values.

Then, the values of the parameters producing the best fit were selected. For the next stage, the minimum number of attractors was found and fixed, present (approximately equivalent) best solutions, obtained with this random search. Then, a grid search for the optimal values of *β* and *α* was performed over a grid of size 30×30, using the logarithmic parameterization described above.

This round of optimization showed that the goodness-of-fit contour lines followed hyperbola-like curves, spanning a broad range of both *β* and *α*, and producing solutions of equivalent quality. Accordingly, the value of attractor strength *α* was fixed at 100, and then, the final search for the best value of the mobility bonus *β* was performed.

### Autocorrelation analysis

To study the distribution of tagged (essential or ancient) genes over the chromosome, and specifically to study the large-distance dependencies between tags, there was used an empirical autocorrelation function, defined as follows. Like previously, the chromosome is modeled as a circular sequence (*x*_1_, *x*_2_, …, *x_n_*) of 0’s and 1’s. For a given nonnegative integer lag *t* < *n*, consider a circularly shifted sequence (*y*_1_, *y*_2_, …, *y_n_*), where *y_i_* = *x*_i+t_ if *i* + *t* ≤ *n* and *y_i_* = *x*_i+t-n_ otherwise. The value *R*(*t*) at point *t* is taken equal to Pearson correlation coefficient between (*x*_1_, *x*_2_, …, *x_n_*) and (*y*_1_, *y*_2_, …, *y_n_*).

The autocorrelation function reveals the dependency between the tags of the genes that lie at a fixed distance from each other. For example, if for some value of lag *t* the autocorrelation function is close to 1, it means that two genes lying at the distance *t* are likely to be either both tagged or both untagged. That can be interpreted as a presence of clusters of tagged / untagged genes of size larger than *t*.

The typical behavior of *R*(*t*) for real genomes is that it is positive for small values of *t*, then it decreases until reaches negative values, after which it oscillates around zero. Denote the smallest value of *t* for which *R*(*t*) is non-positive as “zero-correlation lag”. At this distance the tags become approximately independent, and therefore it can be considered as a rough measure of characteristic size of the clusters in the chromosome.

Zero-correlation lags for the genomes from the following four sets were calculated: 1) the real genomes; 2) the genomes obtained with the model of segmentation driven by protection of essential genes; 3) the genomes obtained from the model with differential mobility and attractors; 4) the real genomes where contiguous blocks were shuffled randomly while keeping their number and sizes.

In case 2), the parameters of the model were chosen in a range where in equilibrium the model produces genomes whose probability of survival under a single disruption event is close to that of the real genomes (genome sizes and number of essential genes are taken from the real genomes, *r* = 100, *N_e_* varies in range [1, 10^4^] with Q1=30 and Q3=50, *k* is in range [2, 5]). The large value of *r* is selected to ensure that the model achieves equilibrium in a reasonable computation time. We also tried to put *r* = 10 and did not see any notable difference except of the proportional increase of *N_e_* (approximately, Q1 becomes 200 and Q3 becomes 800). We expect that further decrease of *r* with simultaneous increase of *N_e_* will only make evolution slower, but not introduce any changes significant for our conclusions.

In case 3), the parameters of the model were chosen according to the procedure described in the previous section. The number of genomes in each set is equal to the number of the considered real genomes (235).

The results are shown in S9 Figure. Here we demonstrate that, as expected, the first model (protection of essential genes) reproduces a distribution of zero-correlation lags that is similar to that of genomes with randomly permuted blocks, whereas real genomes have significantly larger zero-correlation lags. For most of the real genomes, it was impossible to obtain a genome with such a large zero-correlation lag from model 1 for any reasonable values of the parameters. This holds for any k in the range of [2, 5]. By contrast, the distribution of zero-correlation lags for model 2 was similar to that of the real genomes. The correlation coefficient between logarithms of zero-correlation lags of the real and fitted genomes from model 2 was 0.65.

## Data availability

All data and code generated in this work are available for download at

https://zenodo.org/doi/10.5281/zenodo.10963238

## Results

### Classification of genes by evolutionary age within genomes of a prokaryotic genus

In order to classify bacterial and archaeal proteins by evolutionary conservation, all protein sequences encoded in complete genomes of the same genus were clustered with the sequence similarity threshold of 0.5 (see Methods for details). Protein cluster members were mapped back to the individual genomes and the fraction of genomes encoding at least one cluster member was calculated across the genus for each cluster. Analysis of the distribution of this fraction suggested partitioning all protein-coding genes into three classes, those with the high, medium, and low frequency of occurrence within the respective bacterial or archaeal genus. The specific thresholds were selected to yield the average fractions of high- and low-frequency genes as close to 1/3 of a genome gene complement within a genus as possible (S1 Figure).

Genes that are common across a clade likely originate from the genome of the last common ancestor of that clade, whereas the rare genes most likely are recent acquisition. Thus, the classification of genes by frequency of occurrence within a genus corresponds to the classification of genes by evolutionary age relative to the history of the genus. We will refer to the highest-frequency group of genes as “ancient” and the lowest-frequency group as “young” while keeping in mind the inherently rough nature of such characterizations.

### Sub-chromosome-scale distribution of genes by evolutionary age

First, we selected a representative genome within each genus, namely, the genome in which the individual distributions of ancient and young genes were most similar to the genus-wide average (see Methods). Then, we mapped these genes onto the largest (usually, the only) partition of the representative genome and examined their distribution along that chromosome.

In an overwhelming majority of bacteria and archaea, both ancient and young genes displayed a highly non-uniform genomic distribution. Most of the chromosomes contained long segments that were significantly enriched in either ancient or young genes (Fig 1, S1 Table); χ^2^ test was used to test the hypothesis of the uniform density of the selected genes, see Methods for details). In each partitioning, the segments enriched and depleted for ancient or young genes alternated. Only *Blattabacterium*, *Borreliella* and *Sulcia*, all intracellular parasitic or symbiotic bacteria with small genomes, did not show this trend. Typical, “ancient” genomic segments consisted of 230-360 genes and were enriched by a factor of 2.2-4.0 relative to segments depleted of ancient genes. The segments enriched in young genes consisted of 120-210 genes, with the enrichment factor typically in the range of 2.6-4.2. The lengths of the enriched segments were largely independent of the chromosome length (Spearman rank correlation *R_S_* of -0.14 and -0.05 for ancient and young genes, respectively), and accordingly, the number of segments strongly correlated with the chromosome size (*R_S_* of 0.75 and 0.74, respectively).

**Fig 1.**
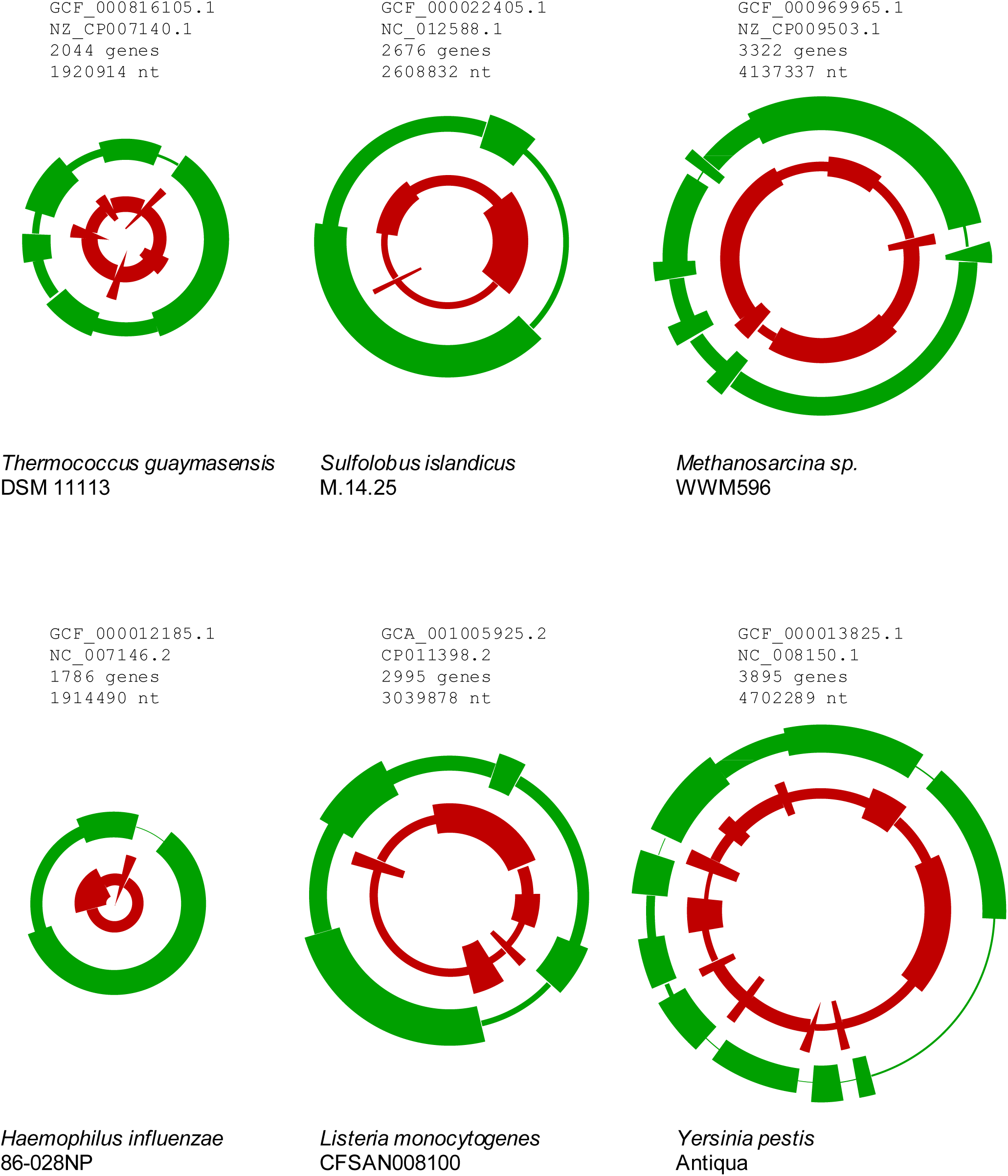
Segmentation of selected archaeal and bacterial chromosomes by gene age. Each circular chromosome is displayed as two concentric rings, showing the average density of ancient (outer ring, green) and young (inner ring, red) genes in chromosome segments that are shown by arcs.

The typical sizes of both the “ancient” and “young” genomic segments, in the (low) hundreds of genes, indicate that they extend well beyond operons and superoperons. Conversely, the sizes of these segments are much less than half of the chromosome size, so that this segmentation clearly does not reflect the leading-lagging strand arrangement or the proximity to the origin of replication. Apparently, evolutionary factors are in action that maintain clustering of genes inherited from the last common ancestor, preventing complete dispersion of these clusters by gene shuffling and insertion of newly acquired genes. Regardless of the exact nature of these factors, they only partially protect the ancient segments from shuffling and gene insertion given that the typical density of ancient genes in such segments was in the range of 0.49-0.67.

Likewise, young genomic segments, although typically shorter than the ancient ones, were also on the scale of 100-200 kbp in length and harbored the same density of young genes (0.50-0.67). In most case, these segments accounted for 50%-70% of all recently acquired genes. This distribution suggests a spatial bias of gene insertions whereby newly acquired genes preferentially (but not exclusively) insert into young genomic segments.

### Distribution of genes at different levels of evolutionary conservation and essential genes in chromosome segments

In an attempt to gain further insight into the driving forces behind the non-uniform distribution of ancient and young genes across prokaryotic chromosomes, we investigated in detail the chromosome segmentation in two well studied model organisms, the bacterium *Escherichia coli* K-12 sub. BW25113 (NZ_CP009273.1) and the archaeon *Sulfolobus islandicus* M.16.4 (NC_012726.1). In addition to genes representing different levels of evolutionary conservation, we explored experimentally identified genes with the lethal knockout phenotype (35,36) that are often used as a proxy for essential genes. Gene essentiality positively correlated with the evolutionary age of genes, but the correlation was not particularly strong, and many essential genes were actually “young” (37) (S2 Table).

In both *Escherichia* and *Sulfolobus*, genes at various levels of conservation and operationally essential genes formed statistically supported segments with similar properties (up to the different fractions of differently tagged genes, Fig 2, S3 Table). None of the alternative evolutionary classifications of genes yielded substantially superior segmentation in terms of the contrast between the enriched and depleted segments compared to the genus-scale operational definition of “ancient” and “young” genes. These findings suggested that the large-scale non-uniform distribution of genes along the chromosome is not driven solely or even mostly by the conservation of the ancestral gene arrangement and/or clustering of evolutionarily conserved genes or functionally important genes as such. Had that been the case, one would expect a stronger segregation of genes that are conserved at greater phylogenetic depths whereas in our analysis, these most ancient genes actually showed a slightly less pronounced segmentation than the genes that were conserved at more shallow levels (Fig 2, S3 Table). However, genes with strong immediate fitness effect (lethal knockout phenotype) segregated even sharper than those conserved at the genus level (although the much smaller number of essential genes makes comparisons difficult). Thus, these findings indicate that gene clustering by age is an evolutionary phenomenon that is manifested at relatively short time scales, likely, reflecting trends in dynamic, comparatively fast processes of chromosome rearrangement.

**Fig 2.**
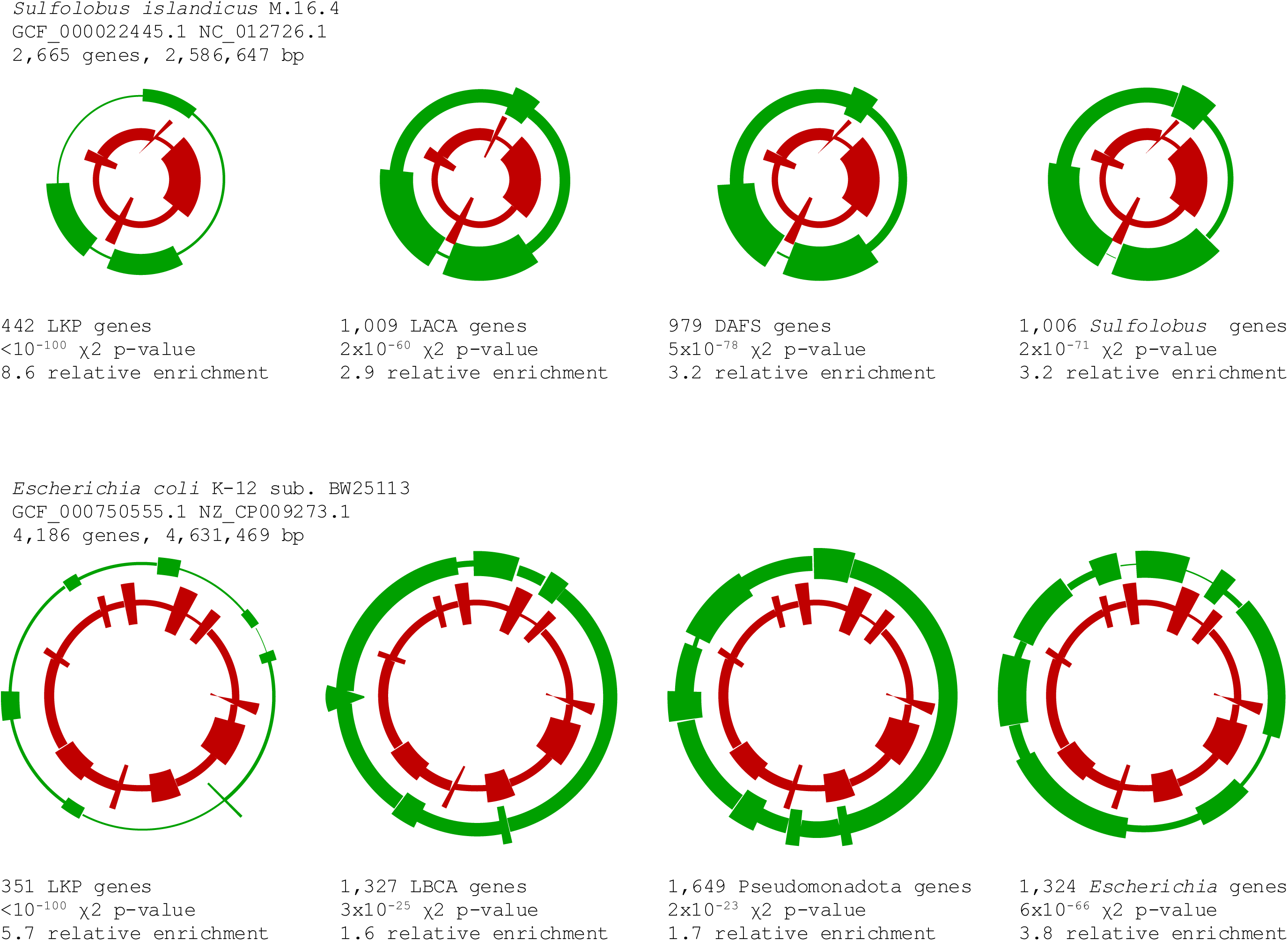
Distribution of genes at different levels of evolutionary conservation and essential genes in the chromosomes of the archaeon *Sulfolobus islandicus* and the bacterium *Escherichia coli*. The inner ring (red) in all panels displays the distribution of young genes. LKP, lethal knockout phenotype genes. LACA, genes likely present in the last archaeal common ancestor. DAFS, genes likely present in the ancestor of *Desulfurococcales*, *Acidilobales*, *Fervidicoccales*, and *Sulfolobales*. LBCA, genes likely present in the last bacterial common ancestor. See Methods for the operational definitions of the ancestral genes.

### Chromosome segmentation and chromatin domains

Both bacterial and archaeal chromosomes have elaborate 3D organization that is maintained through a large part of the cell cycle. The connection of chromatin domains with the properties of the genes located in the corresponding chromosome segments has been demonstrated for the archaea of the genus *Sulfolobus*, where the so-called domain A preferentially harbors highly expressed, conserved genes, whereas domain B contained mostly younger genes expressed at lower levels. The segmentation of *S. islandicus* REY15A chromosome by the density of ancient and young genes observed here roughly corresponded to the chromatin domains A and B (Fig 3A), with domain A, on average, being enriched in ancient genes and domain B in young genes (28). However, a more detailed analysis (S4 Table) showed that this association was far from perfect. Domains A and B were enriched, relative to each other, by factors of 2.19 and 2.57 for ancient and young genes, respectively. For comparison, domain-independent segmentation, based on the observed distribution of ancient and young genes, yielded substantially greater enrichment factors of 3.17 and 4.31 for the ancient and young genes, respectively. The *Sulfolobus* chromatin domains are non-randomly organized relative to the origins of chromosome replication, suggesting that replication polarity and proximity to the origin(s) could contribute to the localization of genes of different ages along the chromosome.

**Fig 3.**
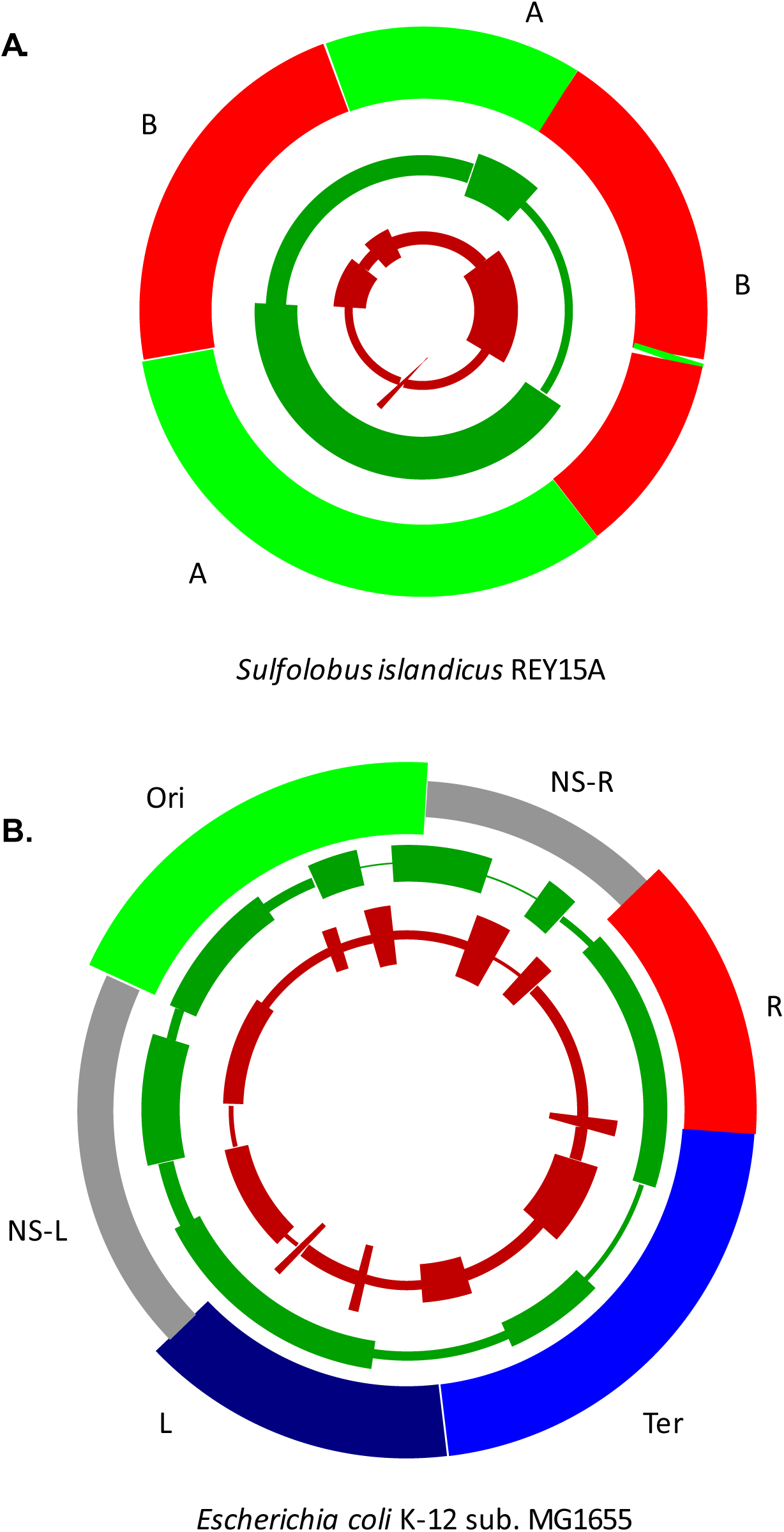
Distribution of ancient and young genes with relation to chromatin domains in the archaeon *Sulfolobus islandicus* and the bacterium *Escherichia coli*. The two chromosome segmentation rings display the distribution of young (red) and ancient (green) genes in *Sulfolobus islandicus* REY15A (panel A) and *Escherichia coli* K-12 sub. MG1655 (panel B). The outer colored rings display chromatin organization domains (A and B for *S. islandicus* (28); Ori, NS-R, R, Ter, L and NS-L for *E. coli* (38).

In contrast, the organization of the chromosome into 6 spatial domains in *E. coli* (38) did not obviously correlate with either the single origin of replication in this bacterium or with the 8 chromosomal segments enriched in ancient genes. Like in *Sulfolobus*, there was a difference in distribution of ancient and young genes between the chromosome domains (the NS-R and TER domains were enriched in young and depleted in ancient genes whereas domains R, L, NS-L and ORI showed the opposite trend), but the signal was weak (relative enrichment factors between the enriched and depleted domains are 1.49 and 1.58 for ancient and young genes, respectively whereas direct chromosome segmentation yielded enrichments of 3.87 and 3.74; Fig 3B, S4 Table).

Taken together, these findings indicated that, even if there was a direct causal connection between chromatin domain organization in *Sulfolobus* and the location of conserved genes on the chromosome, it could not fully explain the segmentation of the chromosome by the gene age. Furthermore, this link was not universal among prokaryotes unlike the segmentation itself.

### Model of evolution of chromosome segmentation driven by protection of essential genes

To explore potential evolutionary factors behind the clustering of ancient as well as essential gene in prokaryotic chromosomes, we then investigated a simple mathematical model of gene order evolution that is driven by fitness effect of multi-gene disruption (12). If adverse events affecting a contiguous region of a chromosome occurred at a non-negligible frequency, then any such event that disrupts an essential gene, is immediately lethal. Because simultaneous disruption of several essential genes is no more lethal than a disruption of a single essential gene, intuitively, grouping essential genes together is beneficial, as it reduces the size of the target for lethal events.

More formally, consider a circular chromosome consisting of *N* genes, *M* of which are labeled as essential and the rest (*N* − *M* genes) as non-essential. Consider an event that occurs at an arbitrary genome location and disrupts a block of *k* adjacent genes. Out of *N* possible genome locations, events in those that include none of the essential genes within a *k*-block, are not lethal (and, as the first approximation, can be considered neutral), whereas the remaining events are lethal. It is easy to estimate the frequencies of such locations for two extreme cases. If all *M* essential genes in a genome form a single block, the fraction *f* = (*M* + *k* − 1)/*N* of events are lethal; if all *M* essential genes are separated by at least *k* − 1 non-essential genes (implying *M* ≤ *N*/*k*), the fraction *f* = *Mk*/*N* of events are lethal which is obviously greater than the fraction for the case of a single block of essential genes.

Let us compare two genome configurations, ***G***_0_ and ***G***_1_, that differ by the arrangement of essential and non-essential genes. Each arrangement is characterized by the relative frequency of lethal disruption events, *f*_0_ and *f*_1_, respectively. Therefore, the impact of the transition from ***G***_0_ to ***G***_1_ can be evaluated by comparing *f*_0_ with *f*_1_. If these frequencies are equal, the transition is neutral, whereas if *f*_1_>*f*_0_, the transition is deleterious, and vice versa. Consider a characteristic rate of gene disruption events of *r* events per genome per generation (with the actual number of occurring events given by the Poisson distribution with the expectation of *r*). An individual will survive if none of these events are lethal; thus, the survival probabilities of ***G***_0_and ***G***_1_ are 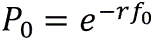 and 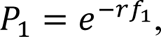, respectively. This allows us to estimate the selection coefficient *s*_0→1_, acting on the configuration ***G***_1_ relative to 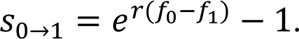 Then, if a gene rearrangement transforms ***G***_0_ into ***G***_1_, the probability of fixation of configuration ***G***_1_ in a population of effective size *N_e_*, is given by the Moran equation (under the low mutation regime) (39): 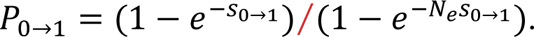

This simple formalism provides for an efficient way to simulate gene order evolution. Starting with a genome configuration ***G***_0_, generate configuration ***G***_1_(e.g. by relocating a randomly selected gene into a randomly selected location), evaluate the frequency of lethal *k*-gene disruptions *f*_1_ for the new configuration, and accept this configuration with probability of *P*_0→1_. A single-gene translocation changes the number of “lethal targets” for *k*-gene disruption events by up to *k* − 1 and is therefore subject to selection of the order magnitude of *r*(*k* − 1)/*N* (assuming *r*(*k* − 1) ≪ *N*). Given the criterion for strong selection, |*s*|*N_e_* ≥ 1, such translocations would be substantially non-neutral if *r*(*k* − 1) ≥ *N*/*N_e_*. Under realistic values of *N* between 10^3^ and 10^4^, *N_e_* ∼10^8^ and *k*>2, an *r* value as low as 10^-3^ is sufficient to render gene translocations far from neutrality.

The numerical simulations demonstrate that this model, provided an appropriate choice of parameters, indeed drives the essential genes to stick together and form a clustered structure. If the selection is strong enough, the number of contiguous blocks of essential genes almost monotonically decreases, and the evolution eventually ends up consolidating all the essential genes into a single, large block (Fig 4a). For weaker selection, this process is more stochastic, and at equilibrium, the number of blocks fluctuates around some intermediate value between perfect clustering and random distribution of essential genes on the genome (Fig 4b-d).

**Fig 4.**
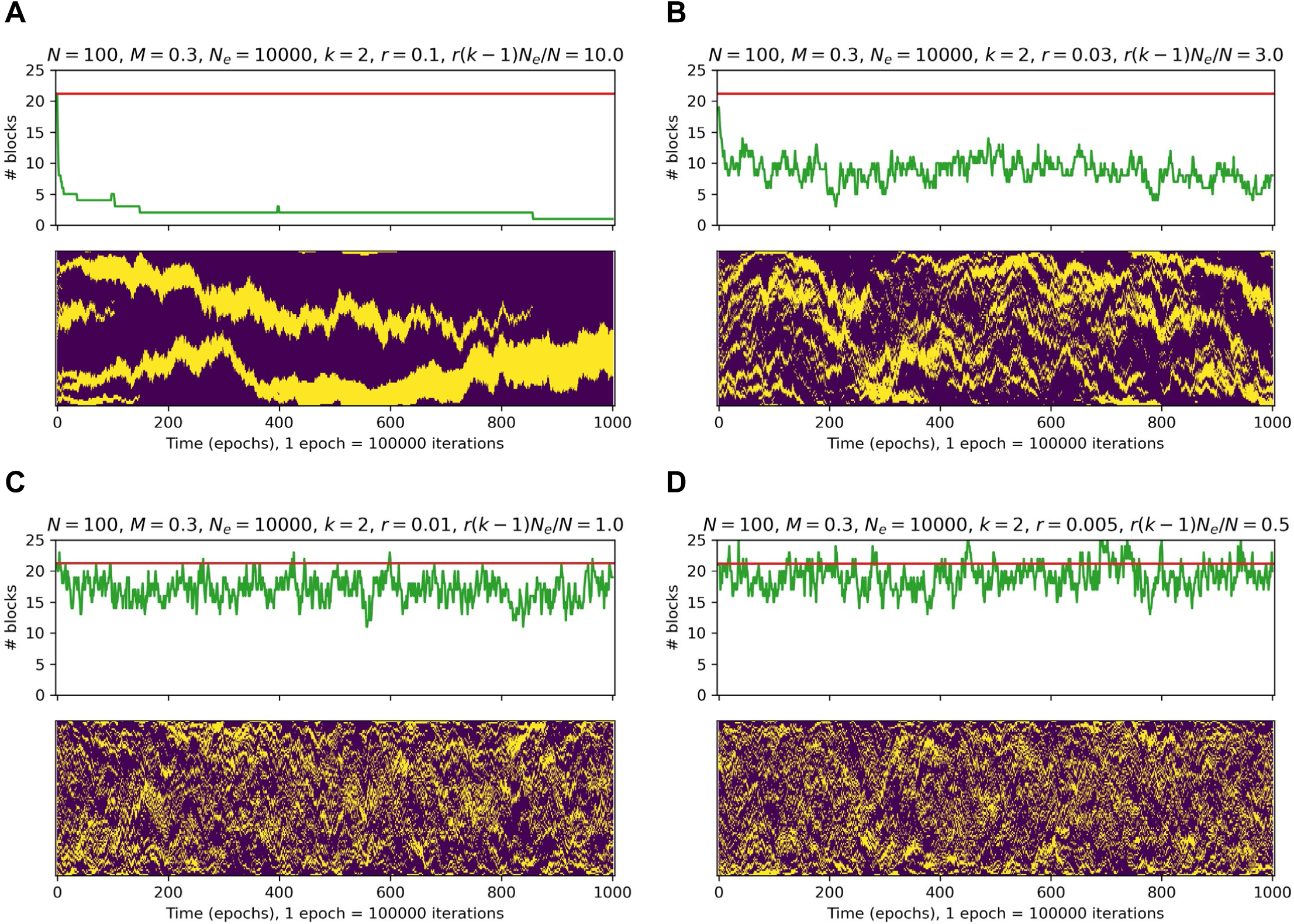
Simulation of gene order evolution under the selection pressure against the effect of multi-gene disruption. On each panel: top, the dynamics of the number of contiguous blocks with evolution time. The horizontal red line shows the mean number of contiguous blocks for a random genome of the given size, with the given number of essential genes. Bottom, evolution of the genome. The genome is represented as a column of colored dots with yellow dots depicting essential genes and magenta dots depicting non-essential genes. The simulated genome state is depicted at every 100000th model step (along the horizontal axis). The model parameters: *N* = 100 (genome size); *M* = 30 (number of essential genes); *N_e_* = 10000 (effective population size); *k* = 2 (number of genes disrupted per event); and different values of *r* (rate of multi-gene disruptions per genome per generation).

However, there was a qualitative difference between essential gene clustering patterns produced by the model and the real data. Specifically, the model results lacked the long-range partitioning of genomes into segments enriched and depleted in essential (ancient) genes that are typical of the real genomes. This difference was most apparent for *k* = 2 where the model did not consistently yield any “fuzzy” segments with the density of essential genes below 1 but significantly higher than that in a randomly shuffled genome. Indeed, the survival probability in this case depends only on the number of solid blocks of essential genes (the lethal target size is equal to the number of essential genes plus the number of solid blocks), and thus, evolution is neutral with respect to the positions of these blocks, provided that they do not merge. This is in stark contrast to the real genomes where we consistently observed segments with the densities of the protected (ancient or essential) genes in the range of 0.5-0.7 (S5 Table and S2 Figure). Numerical simulations confirm these considerations and demonstrate that substantial difference between simulated and real genome also takes place for reasonable values of *k* > 2 (see Methods).

### Genome rearrangement and gene acquisition hotspots

Given that the simplest model of genome evolution driven by selection for minimization of the damage to essential genes failed to capture the salient features of chromosome segmentation by gene age and essentiality, we examined patterns of genome rearrangement in prokaryotes in search of clues to the evolutionary factors behind the observed segmentation. We investigated the short-range evolution of chromosome segmentation (S3 Figure and S4 Figure) and gene order in one archaeal and one bacterial genus by constructing genomic dot plots for pairs of genomes separated by different evolutionary distances and mapping the ancient and young genes onto these plots (S5 Figure and S6 Figure). Examination of these plots showed that genome rearrangements appeared to typically start in a few well-defined hotspots and initially involved acquisition, loss, and translocation of relatively young genes. In agreement with previous findings, many rearrangements involved local segmental inversions (1,14,15). Gradually, as the evolutionary distance between the compared genomes increased, the zones of widespread rearrangement grew and involved more and more ancient genes, until the similarity of the gene order completely deteriorated on the scale beyond (super)operons (S5 Figure and S6 Figure).

The key observation from this analysis was that, despite the loss of the 100 kbp-scale synteny, the segmented structure of the chromosomes with respect to the gene age or essentiality invariably persisted. The preservation of this universal genome segmentation pattern in spite of the disruption of synteny indicated that ancient chromosome segments were not static regions protected from rearrangement and gene insertion, but rather, were dynamic enough to scramble the gene order at the evolutionary distances within a genus. Thus, chromosome segmentation into ancient and young regions appeared to be maintained by biased translocations of ancient genes, whereby operon-scale blocks of ancient genes were likely to end up in ancient gene neighborhoods, and conversely, biased acquisition of young genes, whereby the acquired genes preferentially landed in young neighborhoods.

### Model of gene order evolution with differential gene mobility and translocation attractors

Two observations are salient for understanding the evolution of the chromosome segmentation in prokaryotes: i) at any given time, much of the translocation activity is concentrated in relatively few hotspots on the chromosomes as demonstrated by gene order comparisons at short evolutionary distances, and ii) genes differ substantially in their overall mobility, probably, due to differential fitness effects of synteny disruption, as attested by the preservation of operons over long evolutionary distances. We explored mathematical models of genome evolution, in an attempt to determine whether the pan-prokaryotic chromosome segmentation into ancient and young gene neighborhoods could emerge from these two features of the local evolutionary process without invoking additional factors.

To characterize the (dis)similarity between different patterns (in particular, those observed in real genomes and simulated ones), we employed various metrics, such as absolute difference between the numbers of contiguous gene blocks, mean absolute difference between empirical autocorrelations, and others. Qualitatively, they all lead to the same conclusions. Below, we rely on the recently developed measure, multiscale structural complexity (34), that has been successfully applied in widely different areas (40–43).

The model genomes were represented as circular chromosomes of a fixed size consisting of two classes of genes, those with low and high translocation rates. The fraction of the slow-moving genes is fixed; the relative translocation rate of the faster-moving genes is a parameter. The lower effective rate of translocation implies a fitness penalty on translocation of some genes (selection against translocation with *s* = −3.5 × 10^-4^ reduces the fixation rate by a factor of 10, compared to neutral translocation, in a population with *N_e_* = 10^4^). The model assumes the existence of hotspots (translocation attractors), with the translocation into the vicinity of such an attractor being more likely than translocation elsewhere along the chromosome. The number of attractors, the strength of an attractor (that is, the rate of translocation into a position adjacent to the attractor relative to the rest of the genome) and the size of the attraction basin (how the strength of attraction decays with the distance from the attractor) are the model parameters (although the size of the attraction basin was found to be relatively unimportant and was fixed in most of the simulations).

More formally, consider a circular chromosome consisting of *N* genes. The genes are labeled either as low (*M* genes) or high (*N* − *M* genes) mobility. A low-mobility gene has the relative translocation rate *m_i_* = 1; a high-mobility gene has the relative translocation rate *m_i_* = 1 + *β* (where *β* ≥ 0 represents the mobility factor). A gene is selected for a single-gene translocation event with the probability 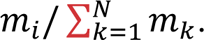. Additionally, *a* out of *N* − *M* high-mobility genes (1 ≤ *a* ≪ *N* − *M*) are labeled as attractors. Each of *N* intergenic regions (IR) can potentially serve as the destination of a translocation. The probability of the *i*-th IR to attract a translocating gene depends on 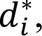, the distance from this IR to the nearest attractor. The relative translocation attraction of an IR is calculated 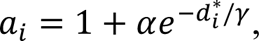, where *α* ≥ 0 represents the attractor strength and *γ* ≥ 0 represents the width of the basin of attraction. An IR is selected to serve as the translocation destination with the probability 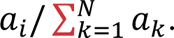 At each discrete time step, a gene is selected for translocation and an IR is selected as the translocation destination with the probabilities defined above.

Within a wide range of parameters (typically, 5-20 attractors per chromosome, with 1-2 orders of magnitude higher translocation rates compared to the rest of the chromosome and with a class of low mobility genes that are 2-20 fold less mobile than the rest), the model reproduced the fuzzily segmented structure of prokaryotic chromosomes, with areas surrounding the attractors enriched in more mobile, young genes and the regions between the attractors enriched in less mobile, ancient genes (Fig 5, S6 Table).

**Fig 5.**
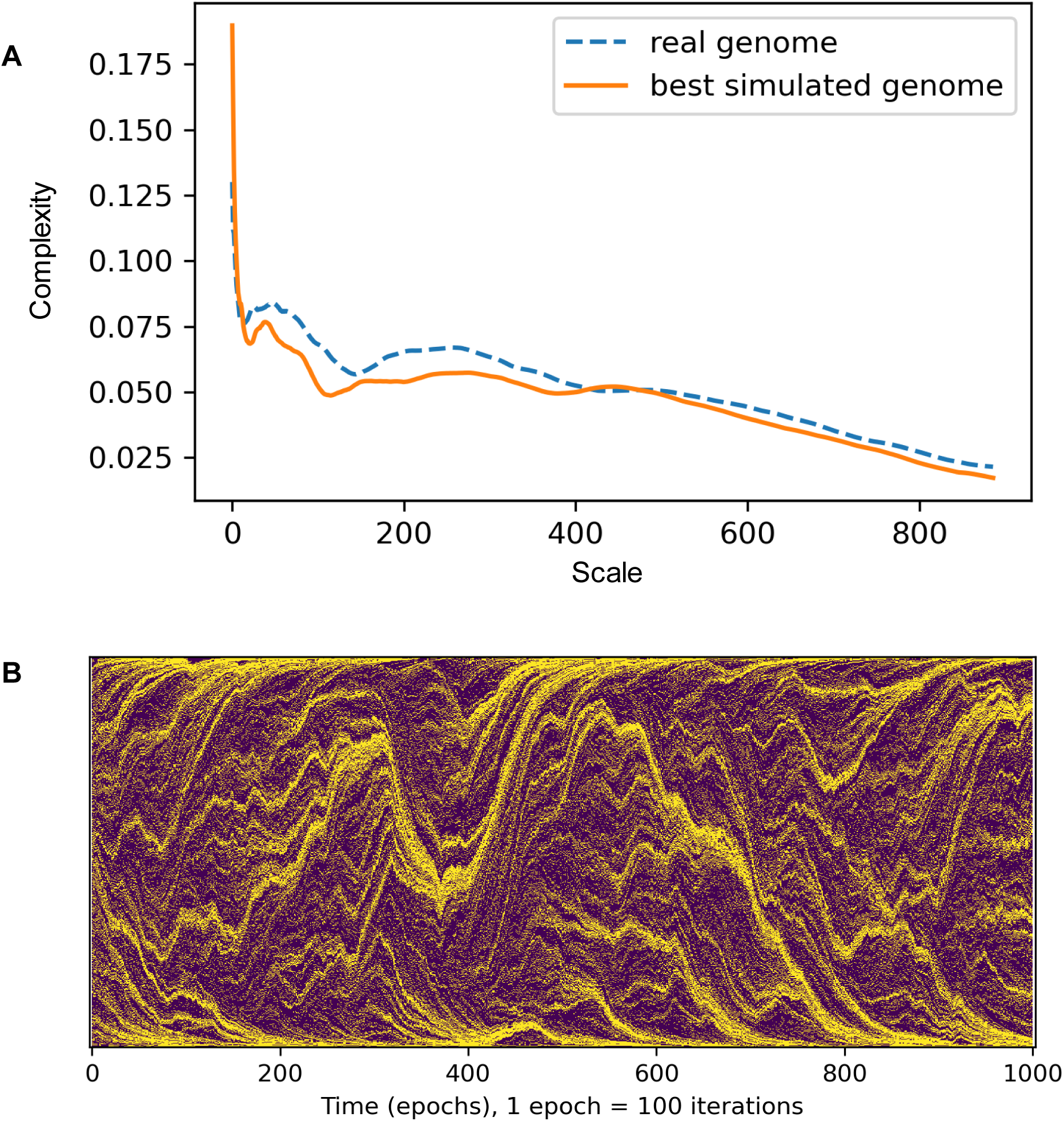
Simulation of gene order evolution under the differential mobility of genes and differential attraction of intergenic regions. A. The comparison of multiscale structural complexity profile between a real and a simulated genome (re.Lactobacillus.NZ_CP062068, dashed blue line; simulation under optimal parameters, orange line). The model parameters: number of attractors: 24, mobility bonus (*β*): 2.49, width of the basin of attraction (*γ*): 1, attractor strength (*α*): 100. B. The genome is represented as a vertical line of colored dots with yellow dots depicting high-mobility genes and magenta dots depicting low-mobility genes. the simulated genome state is depicted at every 100th model step (along the horizontal axis). The model parameters are the same as in panel A.

Both real and simulated genomes display a characteristic multiscale structural complexity profile (see Methods), and thus, the model parameters can be optimized to maximize the similarity between these profiles in simulations. The simulations with the best-fitting models showed significant correlations with the real genomes with respect to some key features (e.g. the density of ancient genes in enriched segments), but for other features, yielded only an overall similarity between the ranges of values (e.g. the mean segment length; S7 Table, S7 Figure). Because the fitting procedure aimed at reproducing the macroscopic properties of the model (reflected in the similarity with the multiscale structural complexity profiles), obtaining a detailed match to the features of real genomes would be unlikely. Nonetheless, despite all the simplifying assumptions, and in contrast to the simple model of genome protection described above, the model reproduced the overall picture of genome segmentation into alternating “fuzzy” segment enriched and depleted in ancient genes with a notable accuracy.

## Discussion

Evolution of prokaryotic genomes is highly dynamic due to extensive HGT, gene loss and genome rearrangements. Although many individual genes are highly conserved through billions of years of evolution, gene order (synteny) is poorly conserved beyond the scale of operons that typically include no more than 5 genes. Therefore, the discovery of universal genome segmentation into mesoscale regions enriched in ancient or, conversely, young genes was unexpected. Our observations indicate that this segmentation actually does not represent evolutionary conservation of gene order, but rather, dynamic persistence of gene localization patterns. Indeed, when ancient genes were defined at a deep phylogenetic level, the observed enrichment in “ancient” genomic segments was even somewhat lower than it was with less highly conserved genes. Thus, the observed segmentation apparently is a dynamic trend that is perpetuated by biased genome rearrangements. This bias is likely to be related to the differential importance of genes for the survival of bacteria and archaea because we observed a particularly pronounced segmentation when genes were classified as essential vs non-essential instead of ancient vs young. Indeed, perhaps, counter-intuitively, gene essentiality in bacteria has been demonstrated to be a poorly conserved, dynamic property, with the sets of essential genes changing substantially on short evolutionary scales (44).

Our attempts to identify a single, simple causative factor behind the observed genome segmentation did not succeed. The characteristic size of the genomic segments enriched in ancient or young genes, in the low hundreds of genes, substantially exceeds the size of even the longest superoperons, ruling out any direct connection to operonic organization. Conversely, these segments are far too short than the half-chromosome scale associated with non-random gene arrangement with respect to the origin of replication. The structural organization of the chromosomes might make a contribution, but it is neither universal nor defining. The simple idea that the segmentation evolves under the pressure for minimizing the probability of the disruption of important genes does not directly apply either because, as shown by our mathematical modeling of genome evolution under this type of selection, the structures it produces differ qualitatively from the real ones with respect to the patterns of the long-range distribution of essential (ancient) genes. The more complex model of genome rearrangements that incorporates two opposing factors, differential gene mobility (a proxy for different fitness effects of gene disruption) and translocation attractors distributed along the chromosome yields a better approximation of the observed segmentation. The translocation attractors do not seem to represent any special physical features of the chromosomes, but rather appear to be regions enriched in young, non-essential genes as well as integrated mobile elements in which insertions are unlikely to be deleterious resulting in preferential attachment of genes gained via HGT. Indeed, it has been shown that in diverse bacterial genomes, the majority of genes recently acquired via HGT concentrate in hotspots that amount to only about 1% of the genome of which many correspond to MGE or various types of genomic islands (45). This model of genome evolution captures the balance between the cohesive force of selection that favors clustering of functionally important genes that tend to be ancient (included in the model as differential gene mobility) and the disruptive process of biased translocation of genome segments that in itself is selectively neutral. However, the existence of translocation attractors might be considered an adaptive feature as well because these chromosome regions can accommodate new genes gained via HGT and hence promote evolutionary innovation and adaptation.

The interplay between differential gene mobility and biased translocation is a clear manifestation of frustration between selective forces that dominate evolution on different scales and appears to be central to all evolutionary processes (46,47). Here, differential gene mobility is a local effect operating at the level of individual genes and operons, whereas biased translocation is a genome-scale phenomenon. In typical physical models the scale separation at pattern formation arises from the peculiarities of the forces, acting on the interacting objects; in our model these forces are not specified in detail, but only through the implications on their effects on gene mobility. It should be noted, however, that this model captures the observed clustering of genes by age or functionality only qualitatively. Thus, additional factors defining the specific features of genome segmentation remain to be identified.

## Author contributions

Y.I.W and E.V.K. designed the study; Y.I.W. and I.V.S. performed research; Y.I.W., I.V.S., K.S.M., M.I.K. and E.V.K. analyzed the data; Y.I.W. and E.V.K. wrote the manuscript that was edited and approved by all authors.

## Acknowledgements

The authors thank Svetlana Karamycheva (NCBI) for expert technical help. I.V.S thanks Askar Iliasov for a fruitful discussion. Y.I.W., K.S.M. and E.V.K. are supported through the Intramural Research Program of the National Institutes of Health (National Library of Medicine). The work of I.V.S. and M.I.K. is supported by the Dutch Research Council (NWO) via the Spinoza Prize of M.I.K.

## Supplementary Figures

**S1 Figure. Classification of “ancient” and “young” genes**

Cumulative distribution function of the protein cluster commonality (fraction of genomes within a genus where members of the cluster are present) is plotted for aggregated genomes of *Sulfolobus* (green line), *Escherichia* (blue line) and *Bacillus* (red line) genera. Horizontal dotted lines indicate the 1/3rd and 2/3rd quantiles, separating the gene complement into three frequency partitions (the least common genes, intermediate commonality genes, and the most common genes). The intersections of these quantile lines with the cumulative distribution curve for a genus *G* correspond to the two quantile thresholds, *Q^L^G* and *Q^H^G*, allowing the classification of protein clusters (and the respective genes) into the least common (“young”) and the most common (“ancient”) categories (displayed for *Sulfolobus*).

**S2 Figure. Segmentation of chromosomes, simulated under the selection against multi-gene disruption**

Comparison of the relative enrichment of enriched segments; the number of segments, average length of enriched segments, fraction of genome in enriched segments, the average density of tagged genes in the enriched segments and the fraction of tagged genes across all enriched segments in simulated and observed chromosomes (y- and x-axis, respectively).

**S3 Figure. Chromosome segmentation within the *Sulfolobus* genus**

Each circular chromosome is displayed as two concentric rings, showing the average density of ancient (outer ring, green) and young (inner ring, red) genes in chromosome segments that are shown by arcs. The tree shows the approximate evolutionary relationships between *Sulfolobus* isolates (48).

**S4 Figure. Chromosome segmentation within the *Lactococcus* genus**

Each circular chromosome is displayed as two concentric rings, showing the average density of ancient (outer ring, green) and young (inner ring, red) genes in chromosome segments that are shown by arcs. The tree shows the approximate evolutionary relationships between *Lactococcus* isolates (48).

**S5 Figure. Evolution of gene order in the *Sulfolobus* genus**

Dots indicate the locations of shared genes (belonging to the same 0.5 amino acid similarity clusters) between two genomes. The color of the dots and the corresponding bars on the horizontal and the vertical edges of the rectangle indicate the estimated evolutionary age of the gene: green, ancient; gray, intermediate; red, young. Panels are arranged in the order of increasing evolutionary distance from *S. islandicus* REY15A (on the horizontal axis in all panels).

**S6 Figure. Evolution of gene order in the *Lactococcus* genus**

Dots indicate the locations of shared genes (belonging to the same 0.5 amino acid similarity clusters) between two genomes. The color of the dots and the corresponding bars on the horizontal and the vertical edges of the rectangle indicate the estimated evolutionary age of the gene: green, ancient; gray, intermediate; red, young. Panels are arranged in the order of increasing evolutionary distance from *L. lactis* subsp. *lactis* S0 (on the horizontal axis in all panels).

**S7 Figure. Segmentation of chromosomes, simulated under the differential mobility of genes and differential attraction of intergenic spacers**

Comparison of the relative enrichment of enriched segments; the number of segments, average length of enriched segments, fraction of genome in enriched segments, the average density of tagged genes in the enriched segments and the fraction of tagged genes across all enriched segments in simulated and observed chromosomes (y- and x-axis, respectively).

**S8 Figure. Cluster commonality thresholds for 175 genera**

The vertical axis shows the cluster commonality range (0 to 1) with lines connecting the lower and the upper thresholds for each genus (*Q^L^G* and *Q^H^G*, respectively), separating the “young”, intermediate, and “ancient” genes.

**S9 Figure. Zero-correlation lags for real, model, and shuffled genomes**

The distributions (kernel density estimation) of logarithms of zero-correlation lags. Model 1 is the “protected essential genes” model. Model 2 is the “differential mobility and attractors” model. Shuffled genomes are obtained from the real ones by random shuffling of contiguous blocks while keeping their number and sizes preserved.

## Supplementary Tables

**S1 Table. Segmentation of archaeal and bacterial chromosomes**

Quantitative characterization of the chromosome partitioning into segments, enriched and depleted with ancient and young genes. Data shown for the representative genomes, selected for each of the 175 genera.

**S2 Table. Breakdown of Lethal Knockout Phenotype genes by evolutionary conservation**

Data shown for genus-level evolutionary conservation in *Sulfolobus islandicus* M.16.4 and *Escherichia coli* K-12 sub. BW25113.

**S3 Table. Segmentation of archaeal and bacterial chromosomes at different levels of evolutionary conservation**

Quantitative characterization of the chromosome partitioning into segments, enriched and depleted with selected genes. Data shown for different levels of evolutionary conservation in *Sulfolobus islandicus* M.16.4 and *Escherichia coli* K-12 sub. BW25113.

**S4 Table. Distribution of ancient genes in chromatin domains**

Mapping of chromatin domains to the chromosomes of *Sulfolobus islandicus* REY15A and *Escherichia coli* K-12 sub. MG1655 and the density of the ancient genes in relatively enriched and depleted segments.

**S5 Table. Simulation of gene order evolution under selection for effect of multi-gene disruption: segmentation of simulated chromosomes**

Quantitative characterization of the simulated chromosome partitioning into segments, enriched and depleted with protected genes. Data is shown for a randomly sampled the state in the model runs with parameters selected for the fraction of lethal 3-gene disruptions to match that in the real chromosomes. Quantitative characteristics of ancient gene segmentation in natural chromosomes are shown for comparison.

**S6 Table. Simulation of gene order evolution under the differential mobility of genes and differential attraction of intergenic spacers: model parameters**

Parameters, maximizing the similarity of the multiscale structural complexity profile between the simulated genome and its prototype. The number of attractors *a* and the differential mobility *β* were optimized; the attractor characteristics (the attraction strength and the basin of attraction) were fixed at *α* = 100 and *γ* = 1 respectively. Results of several runs, producing distinct optimal values of *a* and *β*, are shown for some simulated chromosomes.

**S7 Table. Simulation of gene order evolution under the differential mobility of genes and differential attraction of intergenic spacers: segmentation of simulated chromosomes**

Quantitative characterization of the simulated chromosome partitioning into segments, enriched and depleted with low mobility genes. Data is shown for a randomly sampled the state in multiple independent model runs. Quantitative characteristics of ancient gene segmentation in natural chromosomes are shown for comparison.

