## Supplementary figures and images for "Long range segmentation of prokaryotic genomes by gene age and functionality"

### Supplementary figure 1

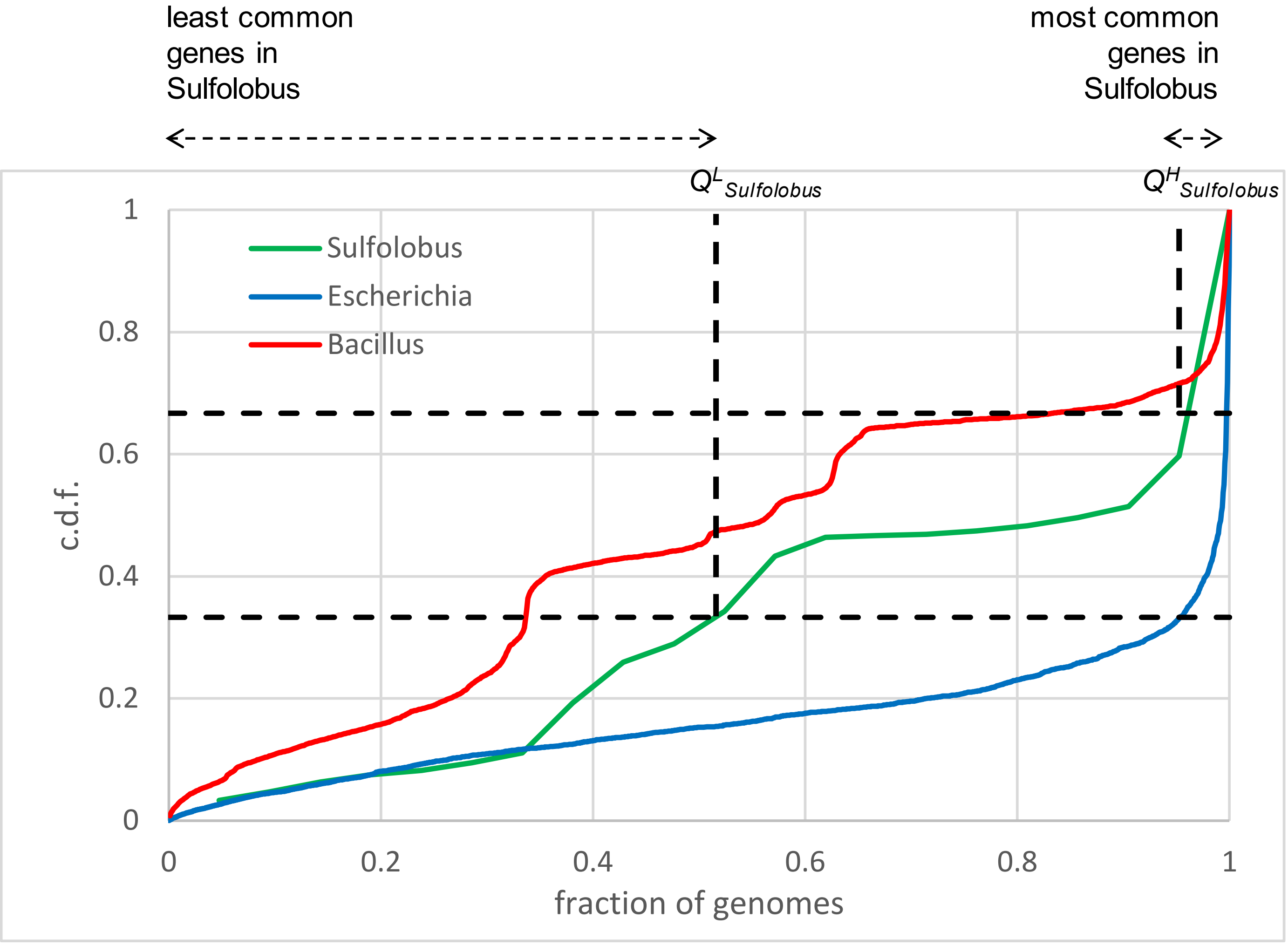

### Supplementary figure 3

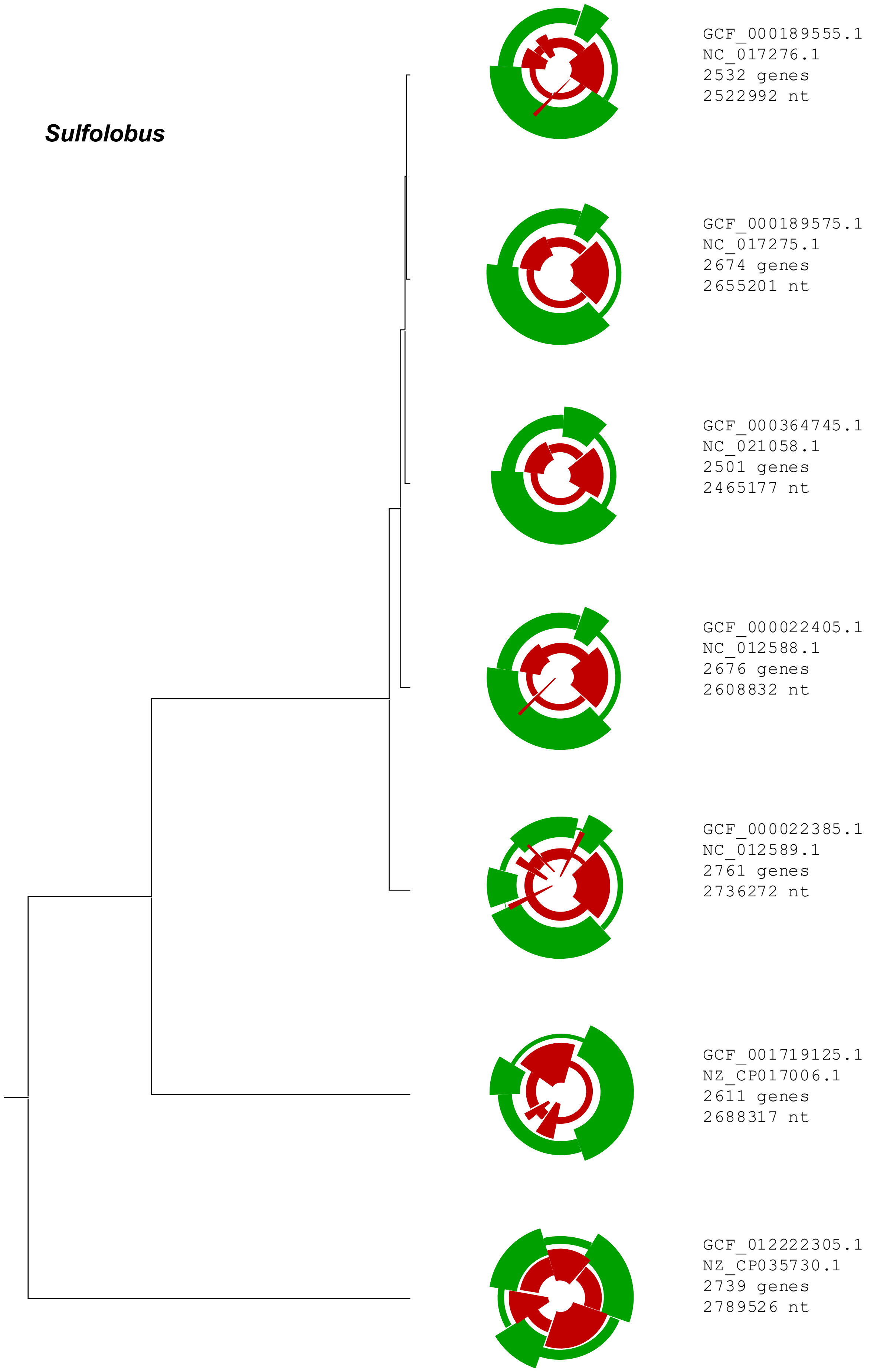

### Supplementary figure 4

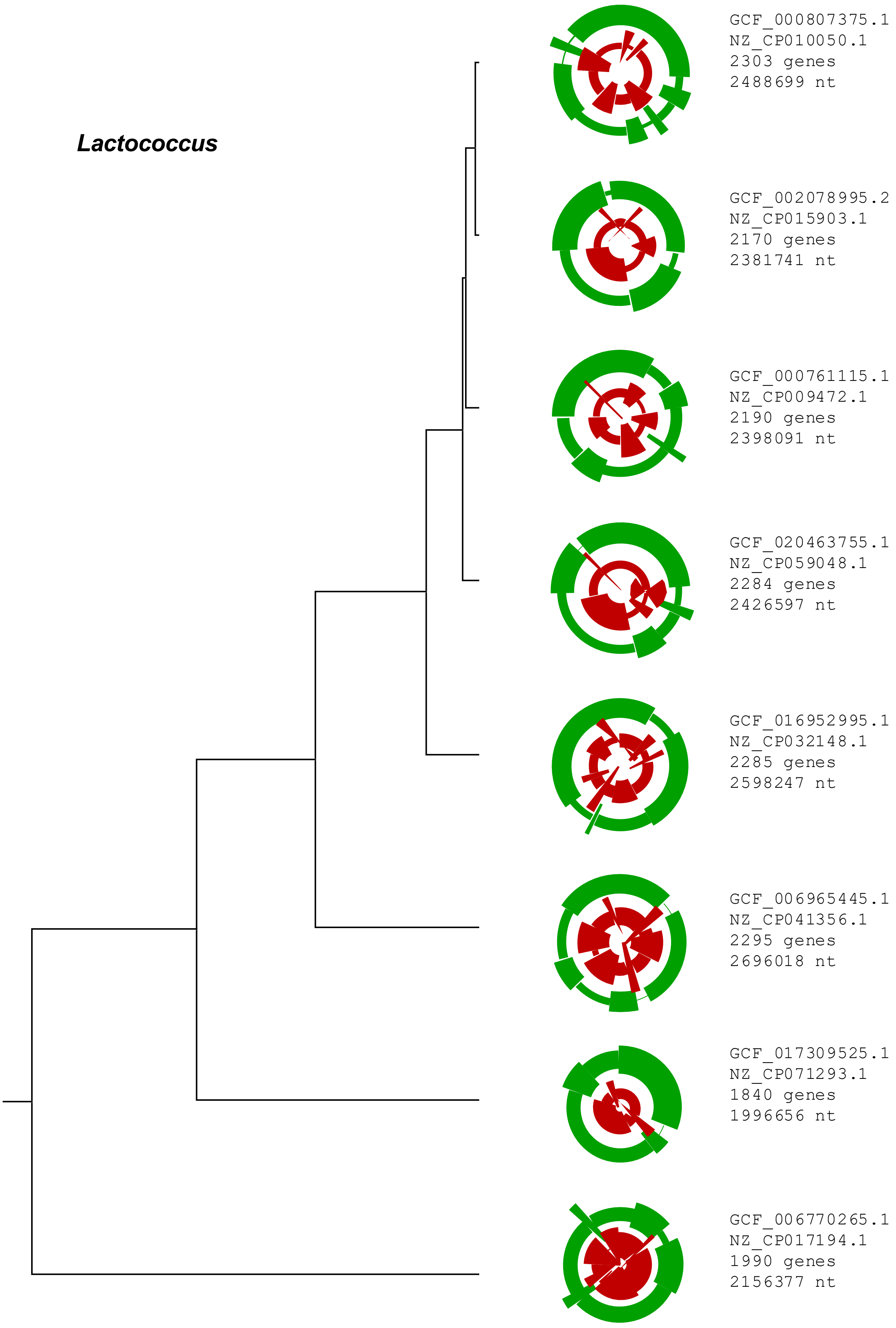

### Supplementary figure 8

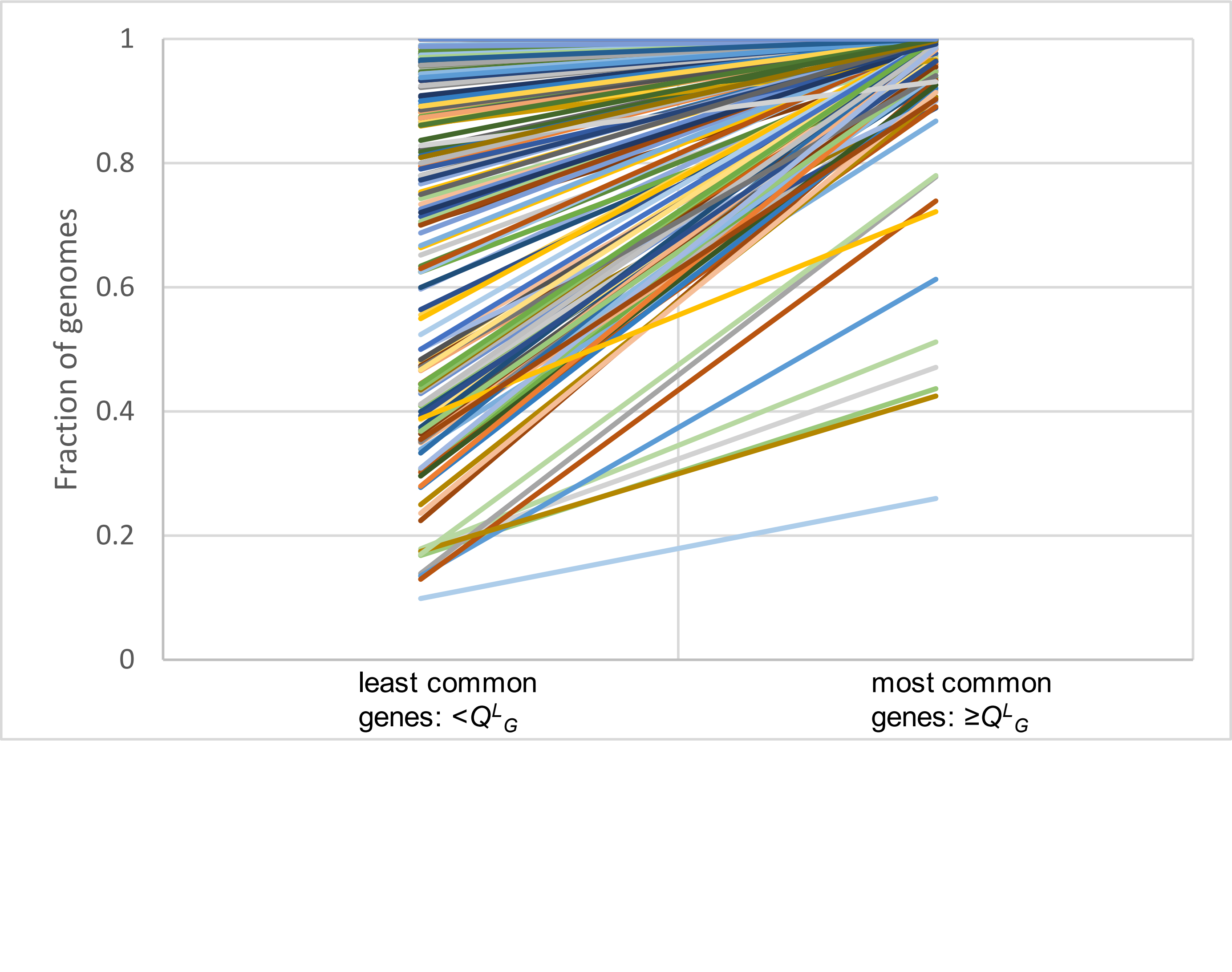

### Supplementary figure 9

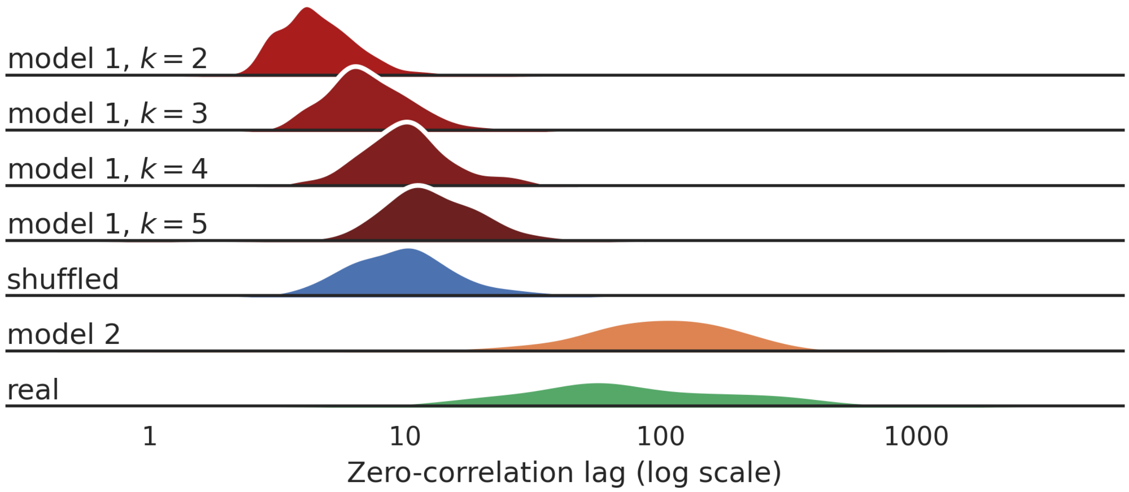
