## Supplementary figure 5 for "Long range segmentation of prokaryotic genomes by gene age and functionality"

### Sulfolobus_islandicus_REY15A

### Sulfolobus_islandicus_HVE104

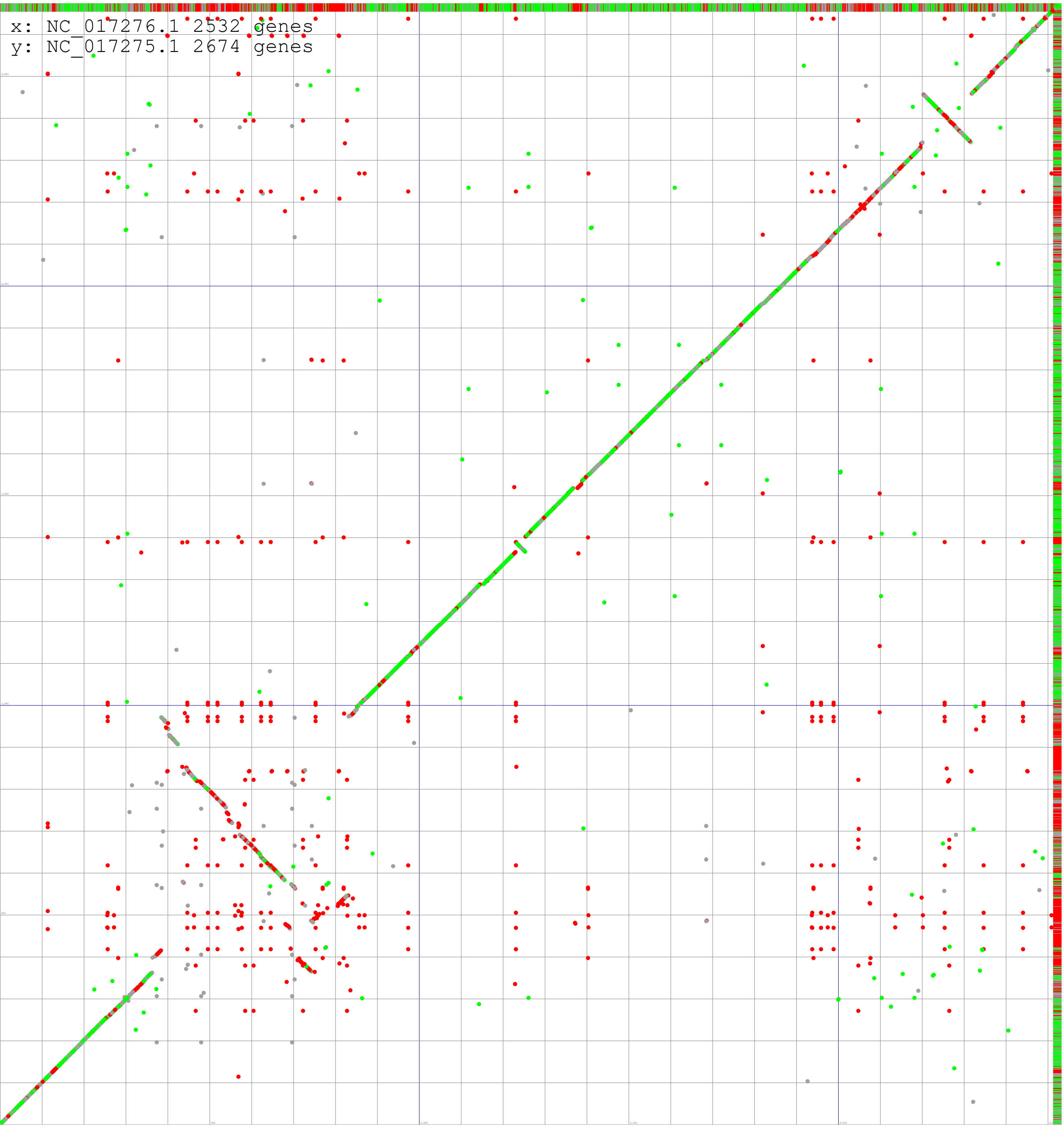

**Sulfolobus_islandicus_REY15A**

**Sulfolobus_islandicus_LAL141**

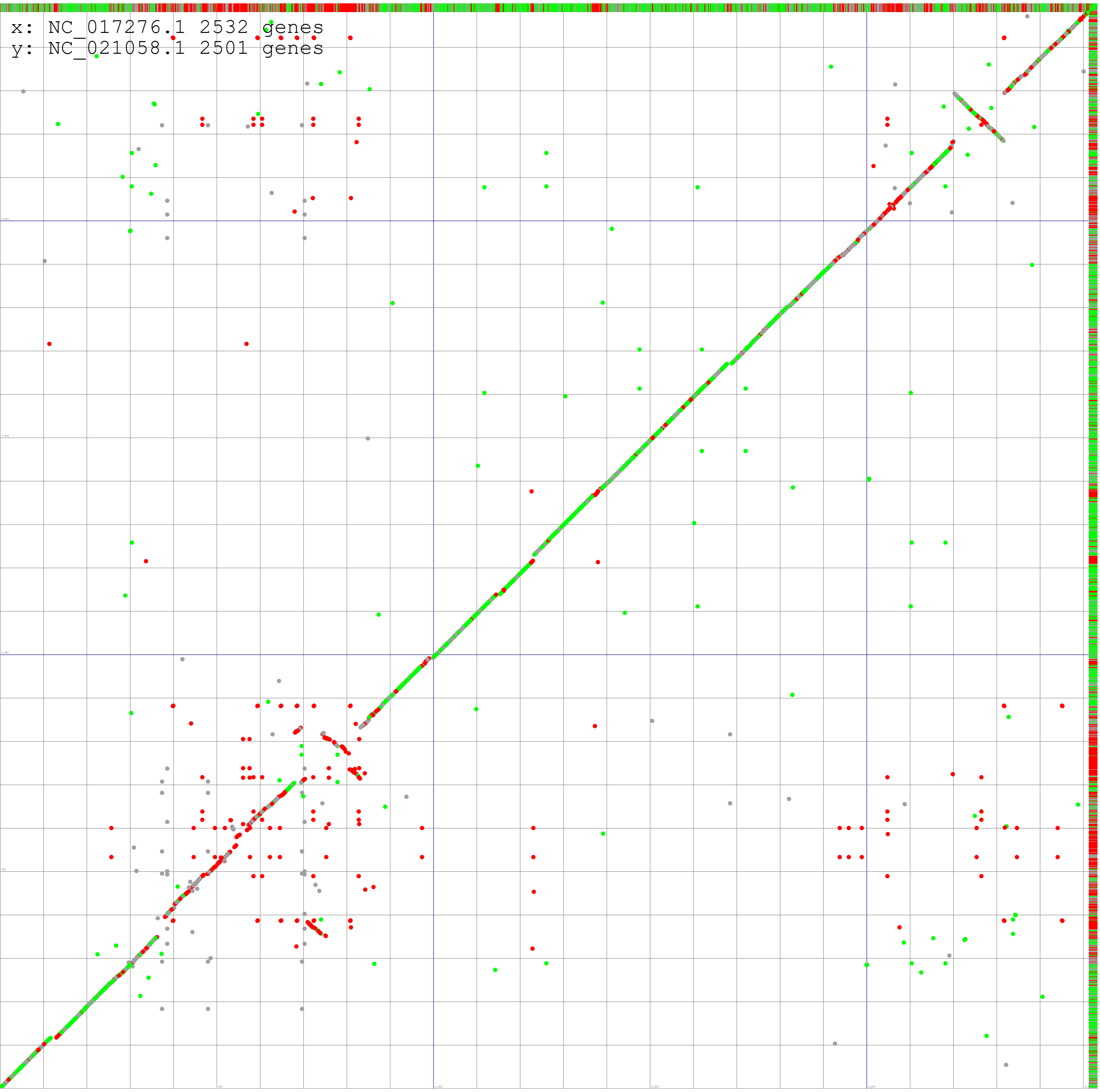

**Sulfolobus_islandicus_REY15A**

**Sulfolobus_islandicus_M.14.25**

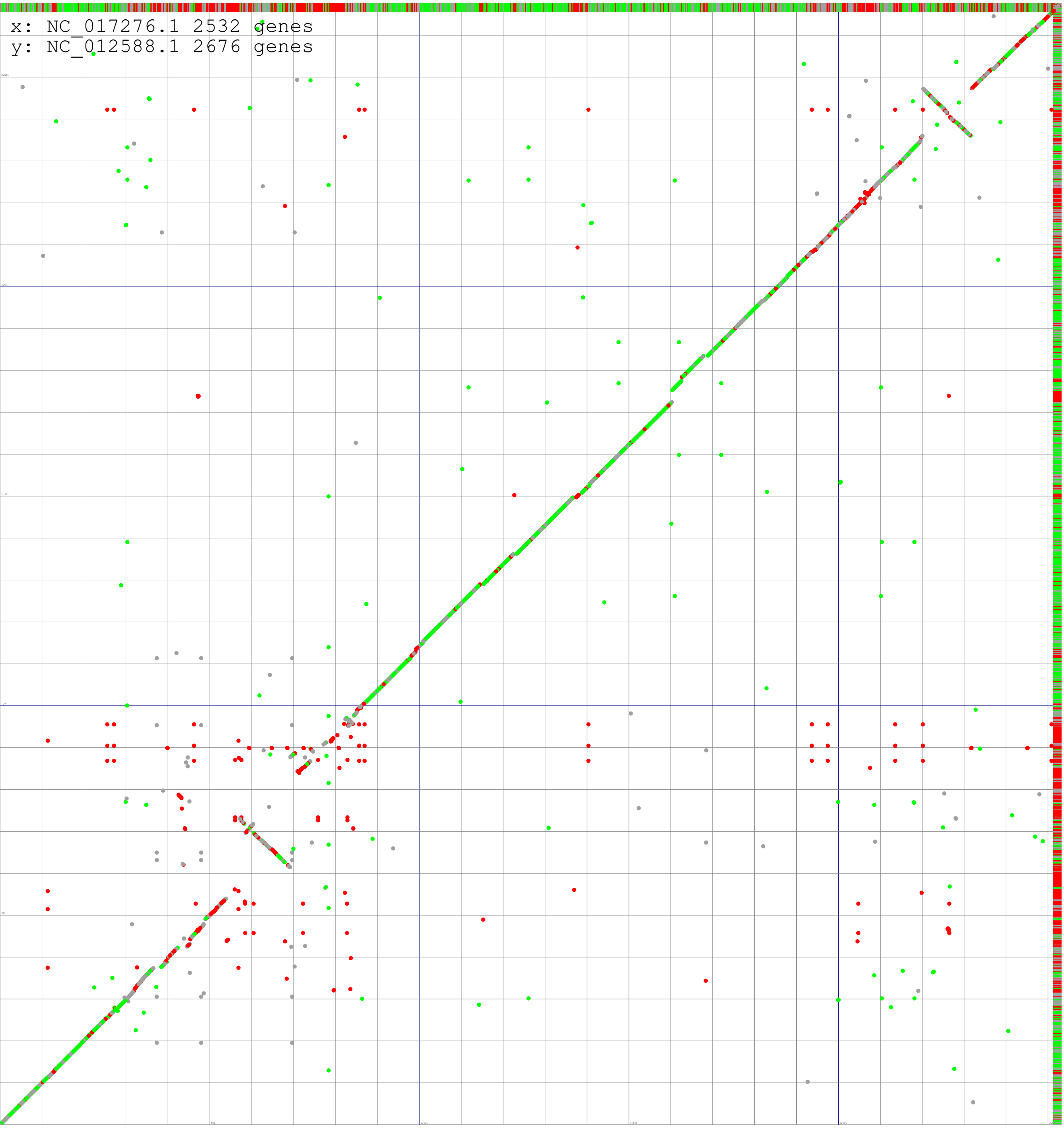

**Sulfolobus_islandicus_REY15A**

**Sulfolobus_islandicus_L.S.2.15**

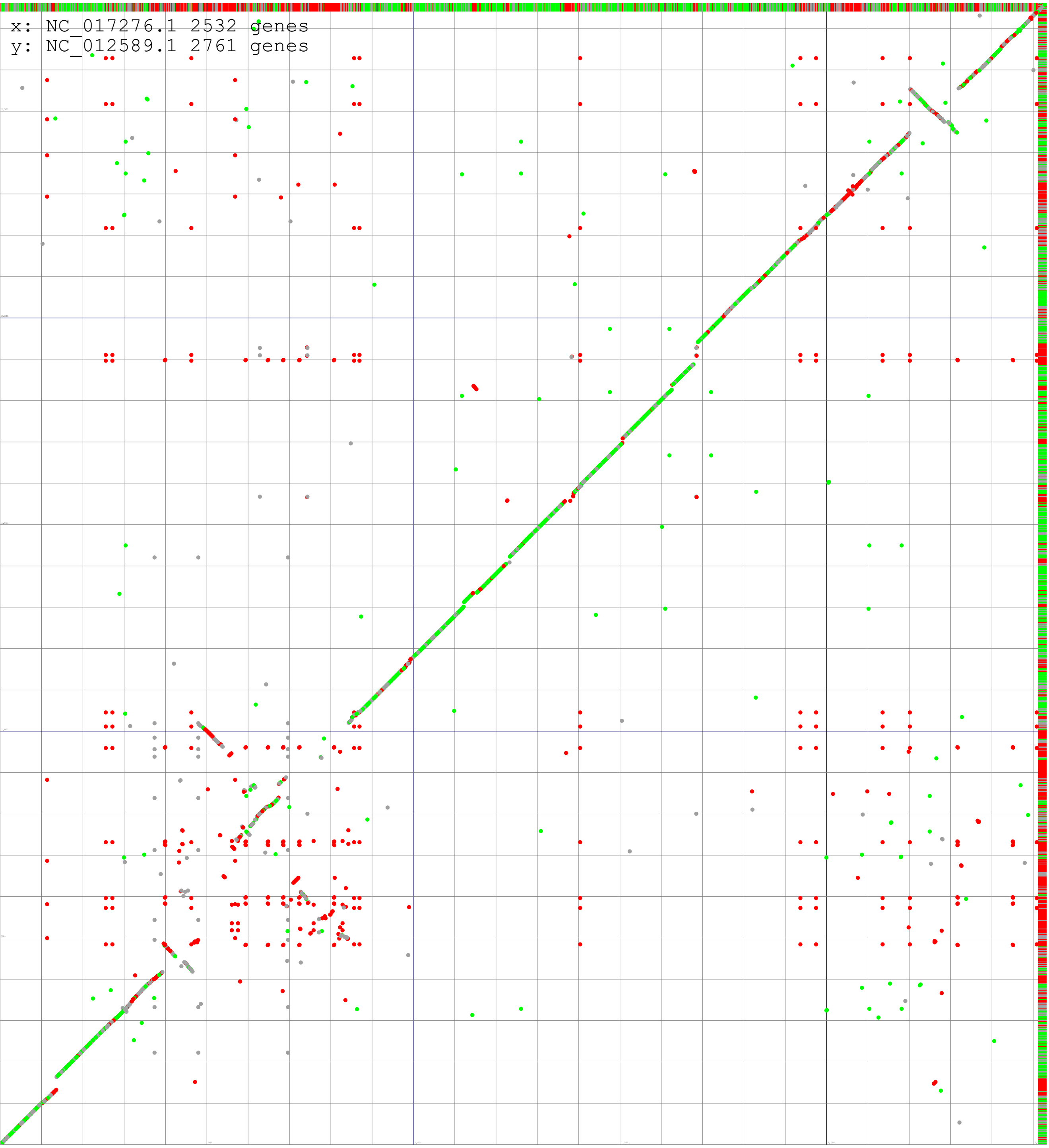

**Sulfolobus_islandicus_REY15A**

**Sulfolobus_A20**

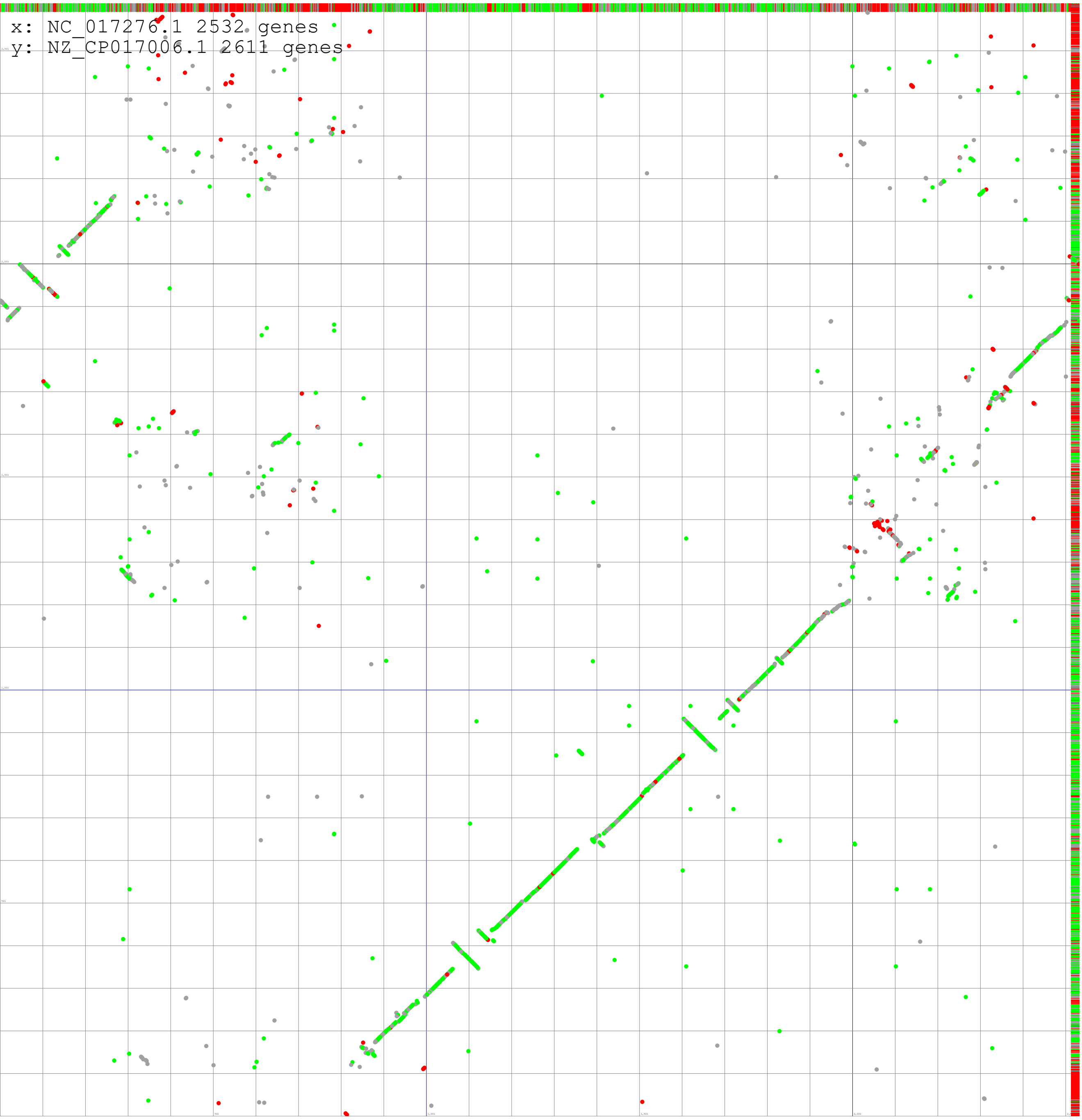

**Sulfolobus_islandicus_REY15A**

**Sulfolobus_S-194**

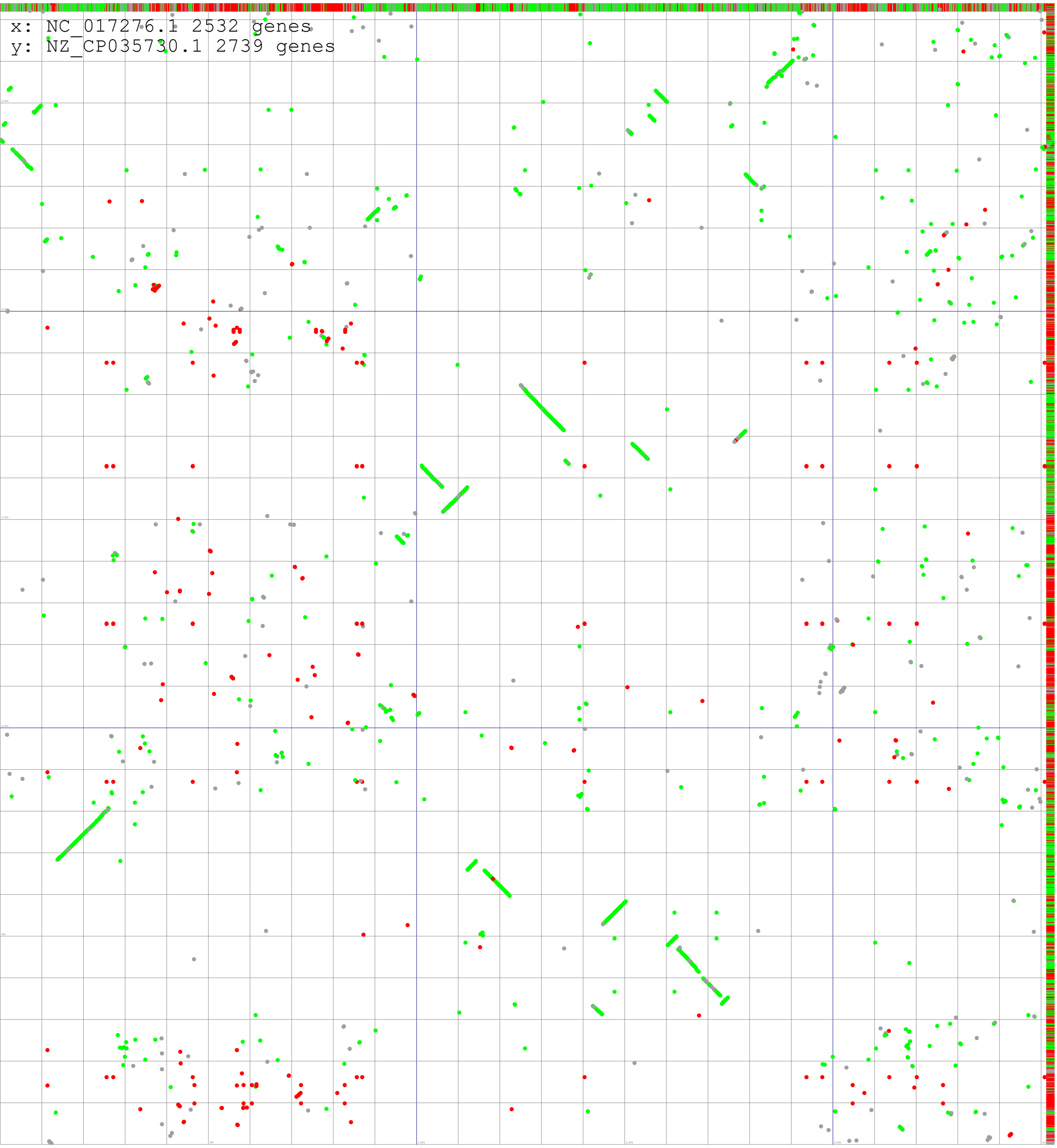
