## Supplementary figure 6 for "Long range segmentation of prokaryotic genomes by gene age and functionality"

**Lactococcus_lactis_sub_lactis_S0**

**Lactococcus_lactis_sub_lactis_UC08**

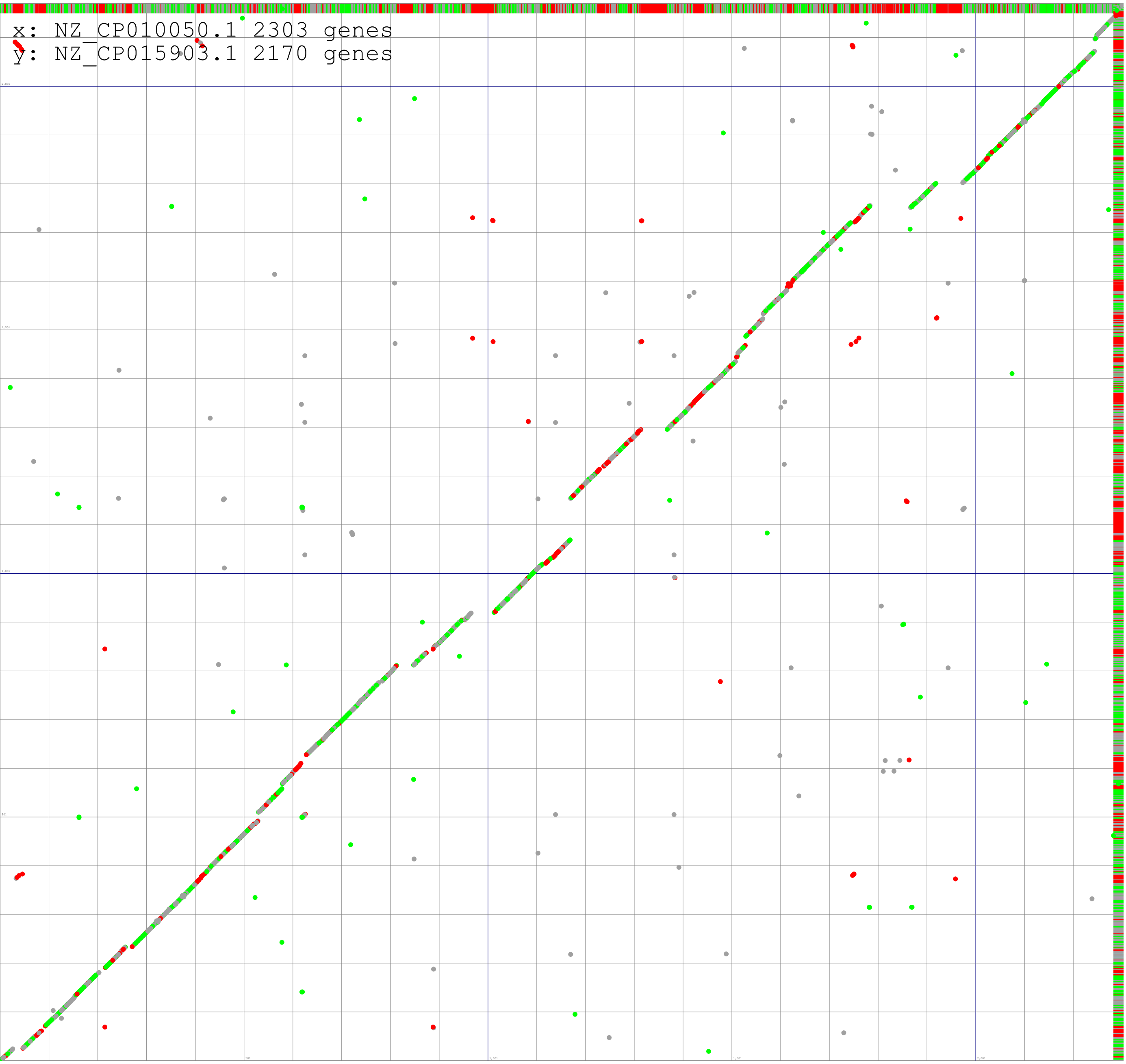

**Lactococcus_lactis_sub_lactis_S0**

**Lactococcus_lactis_AI06**

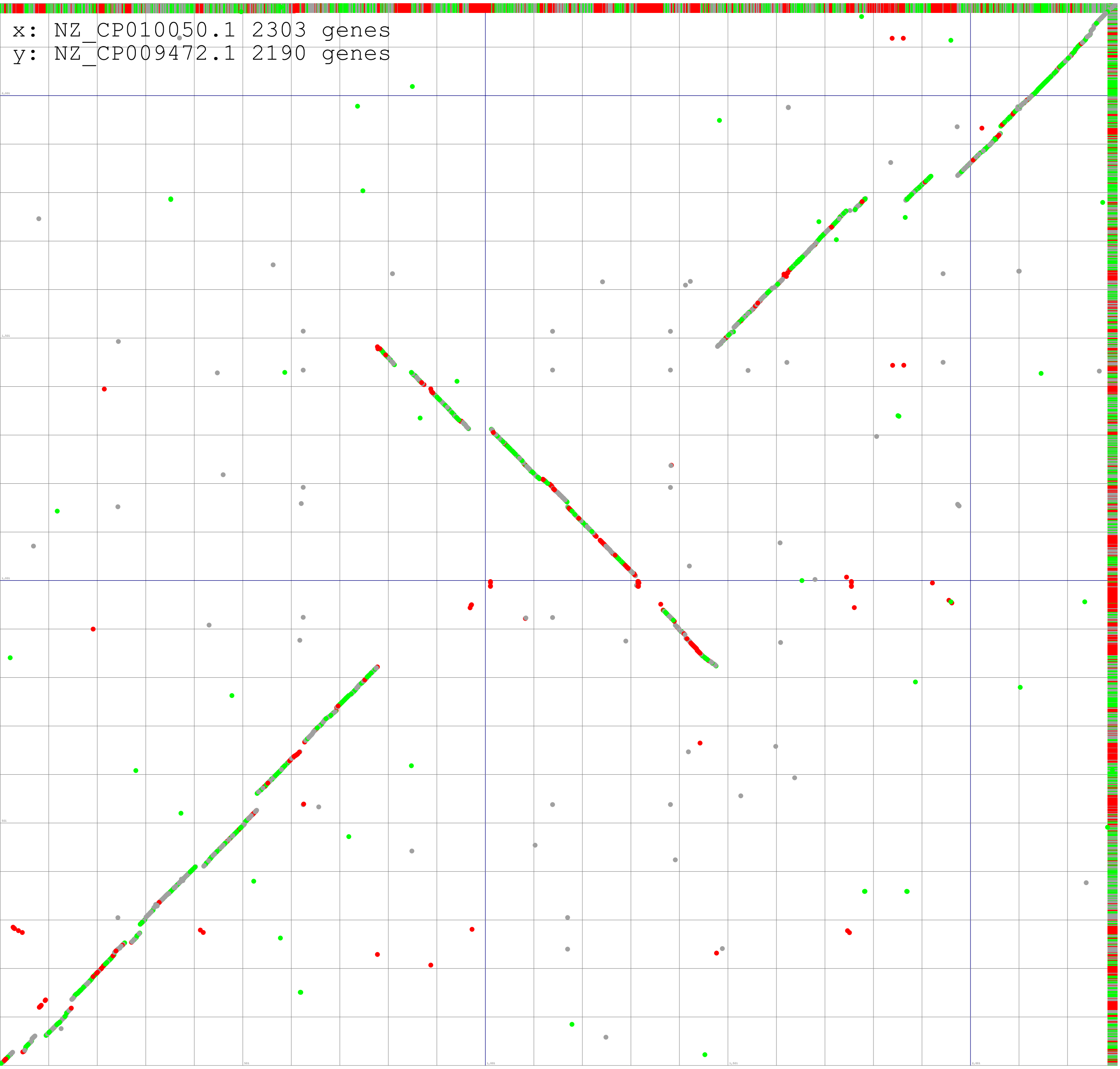

**Lactococcus_lactis_sub_lactis_S0**

**Lactococcus_lactis_LAC460**

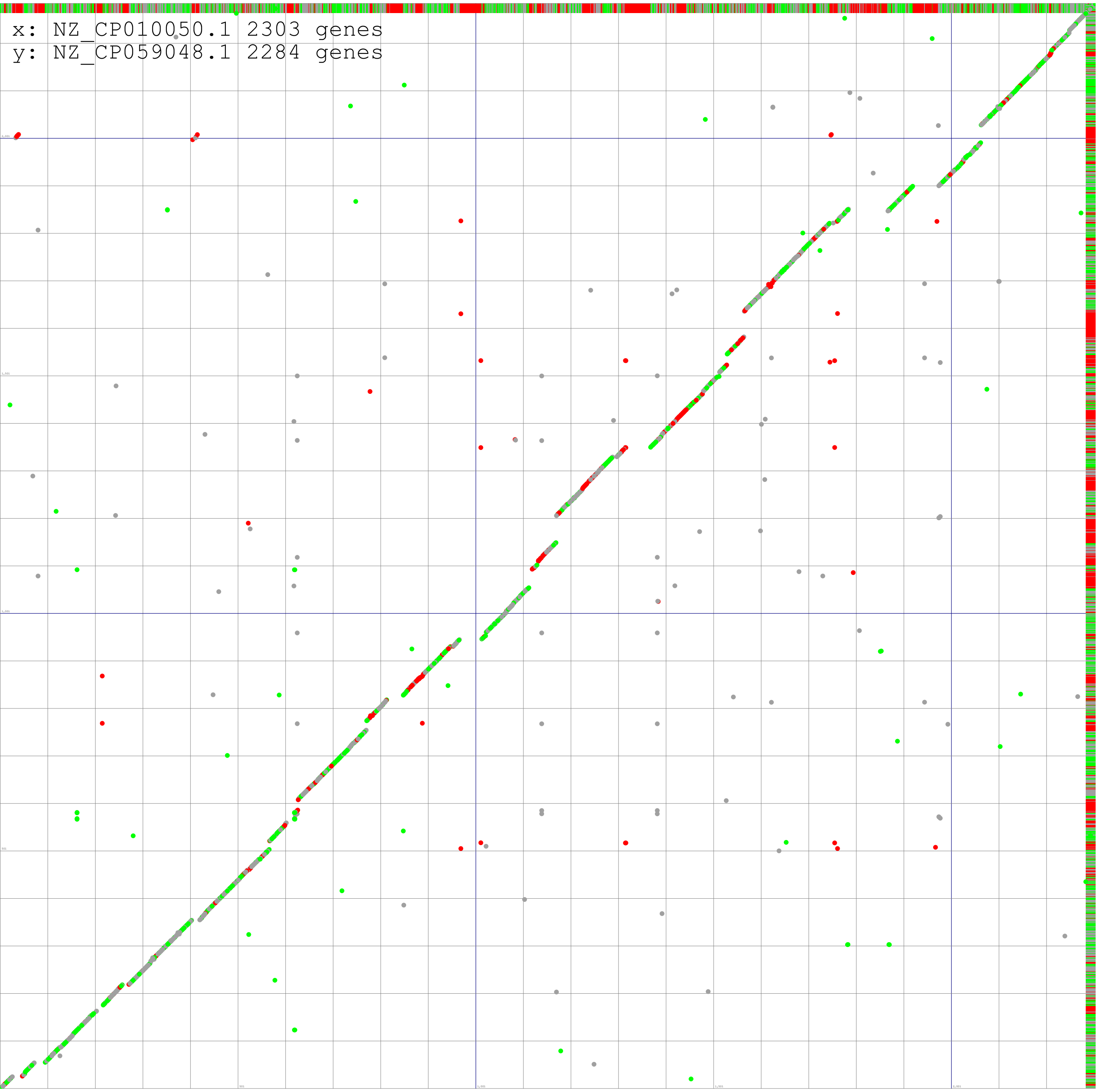

**Lactococcus_lactis_sub_lactis_S0**

**Lactococcus_cremoris_1196**

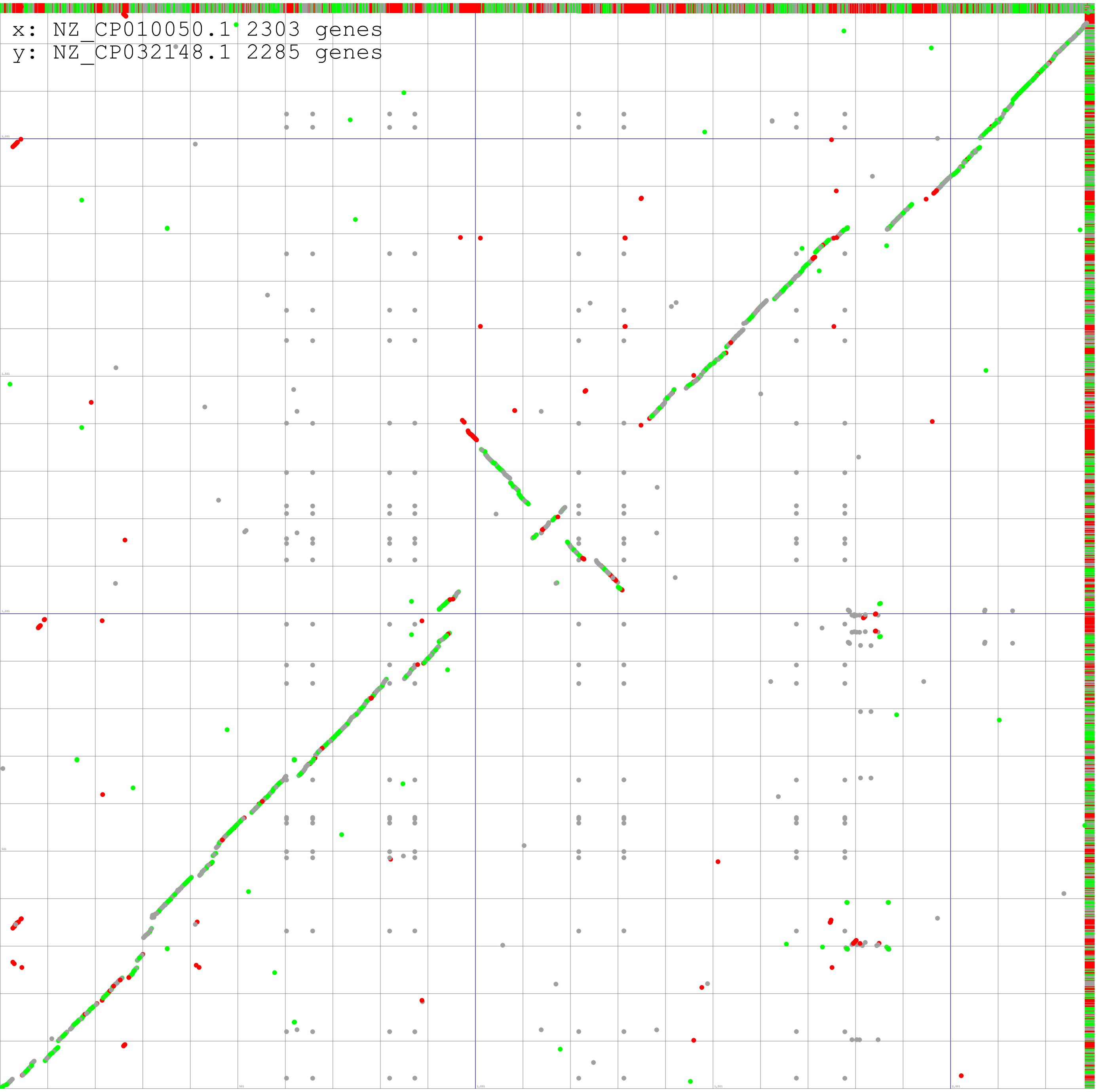

**Lactococcus_lactis_sub_lactis_S0**

**Lactococcus_KACC_19320**

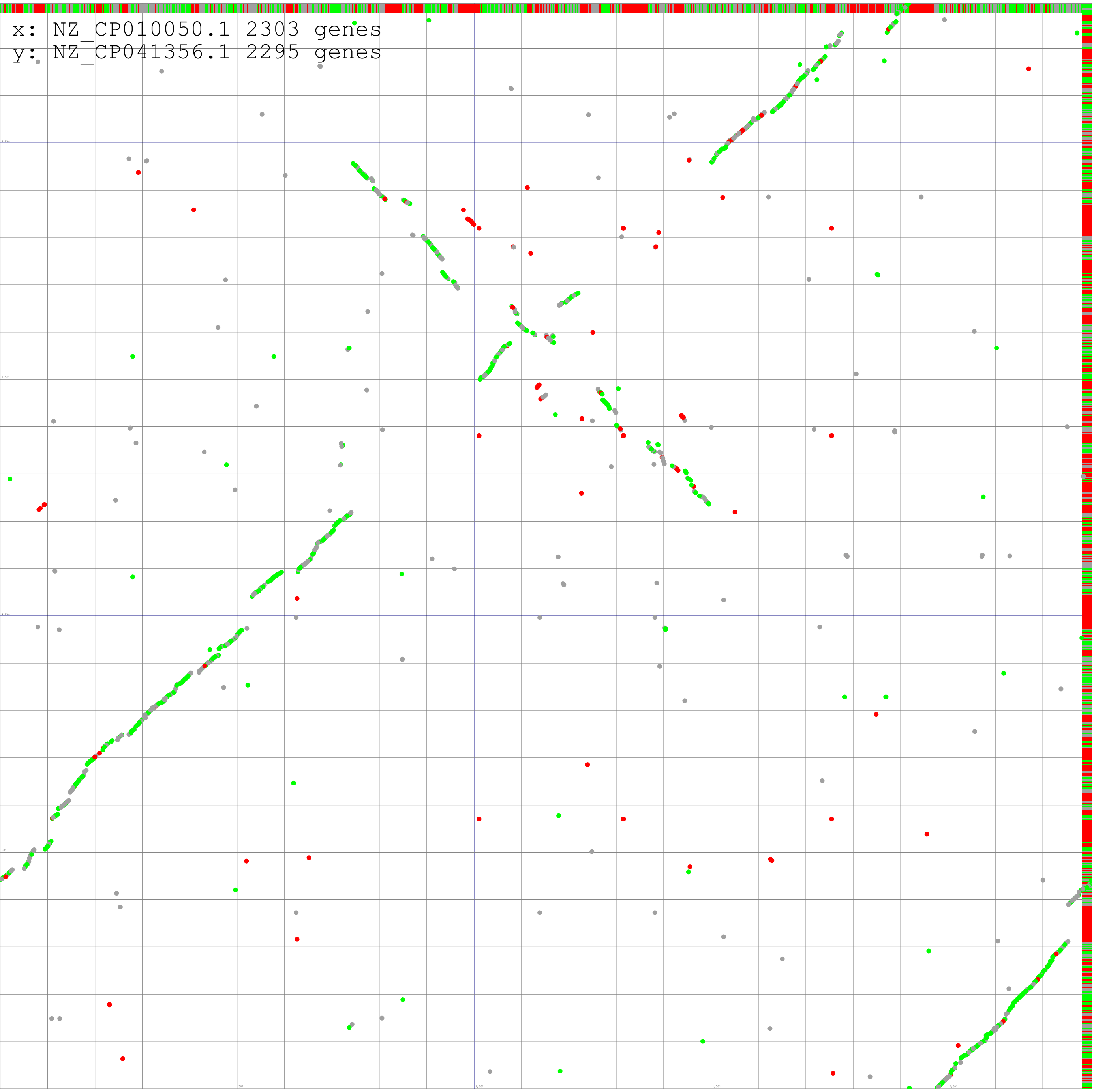

**Lactococcus_lactis_sub_lactis_S0**

**Lactococcus_LG1074**

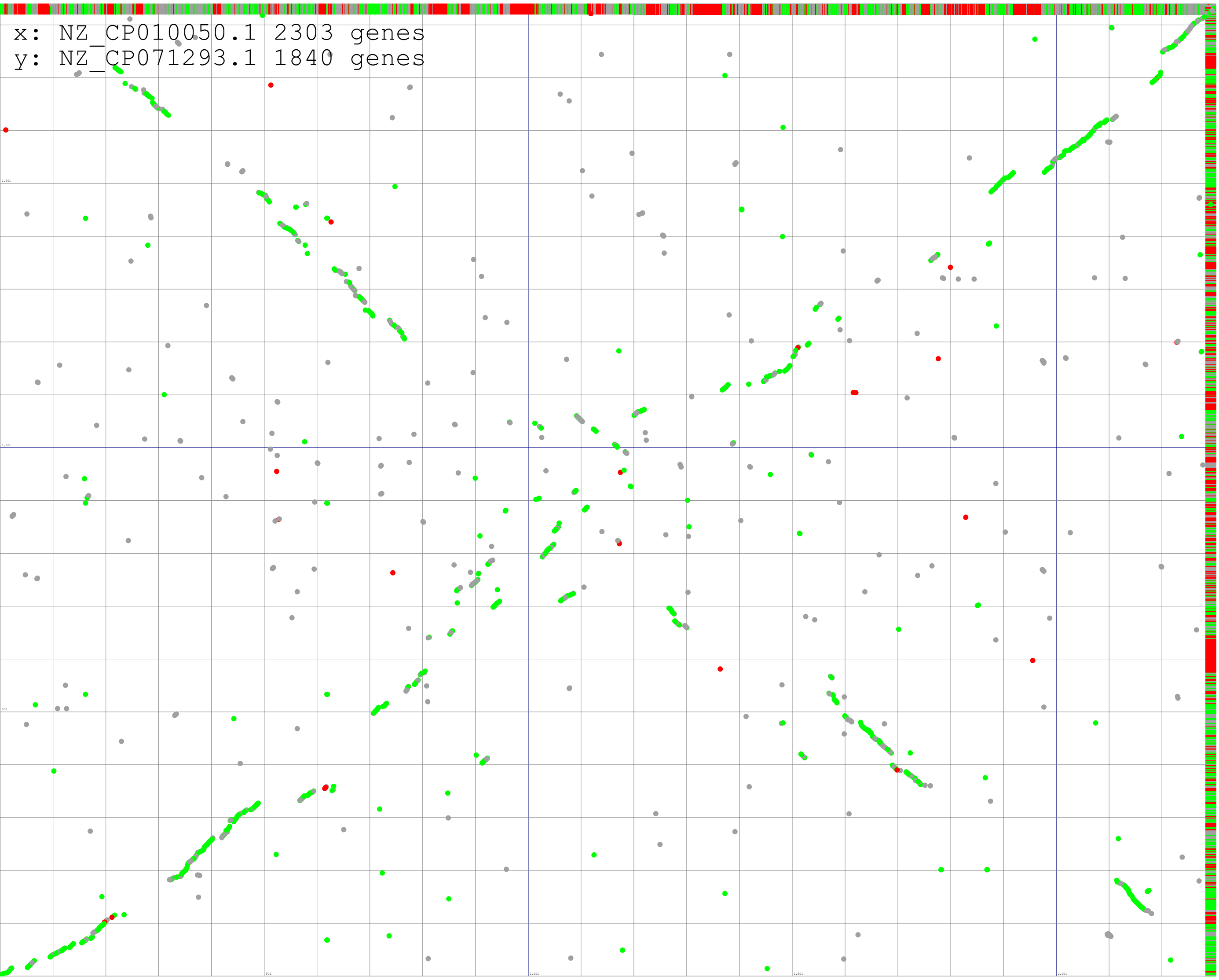

**Lactococcus_lactis_sub_lactis_S0**

**Lactococcus_carnosus_TMW_21612**

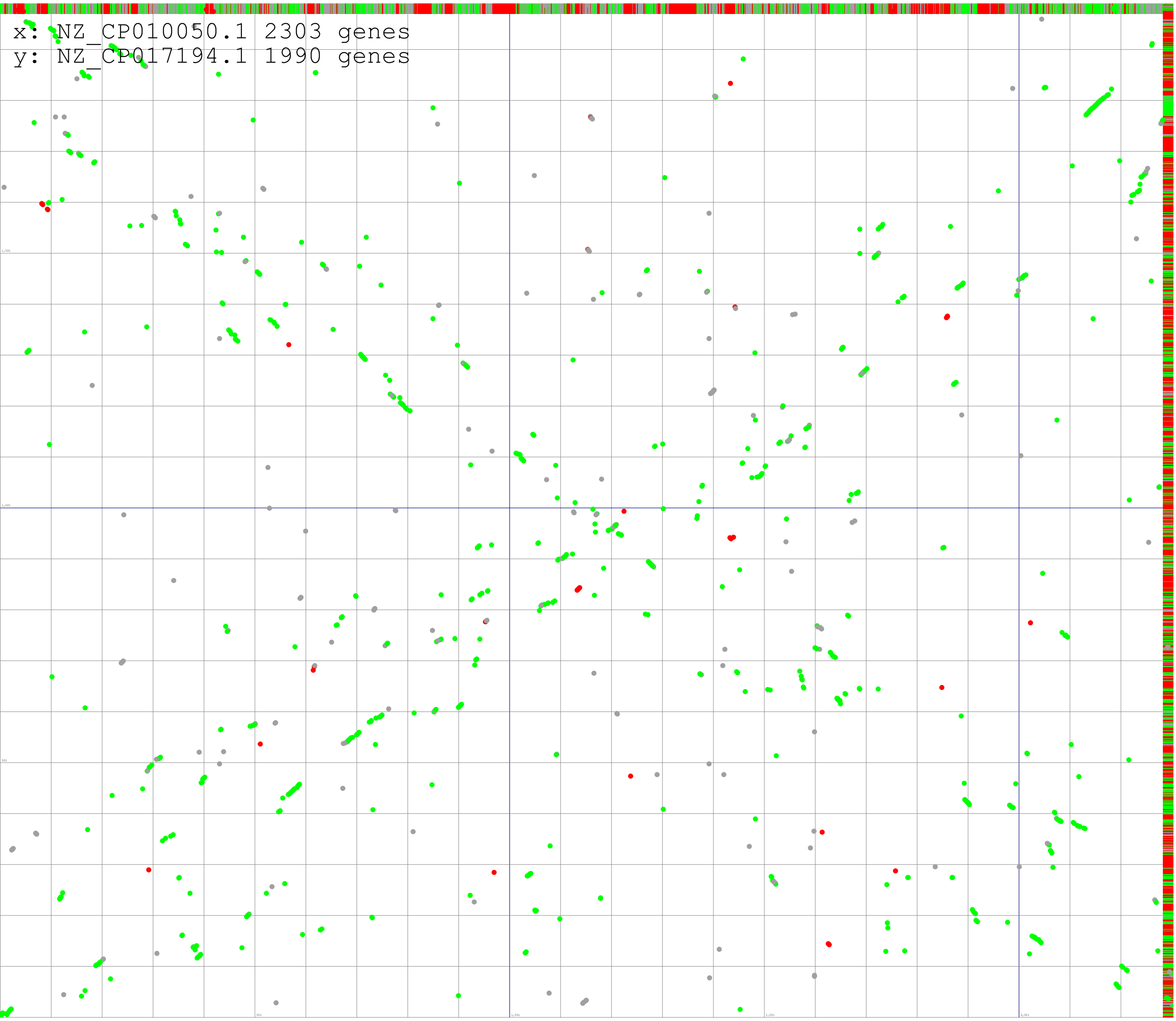
