## Supplementary figure 7 for "Long range segmentation of prokaryotic genomes by gene age and functionality"

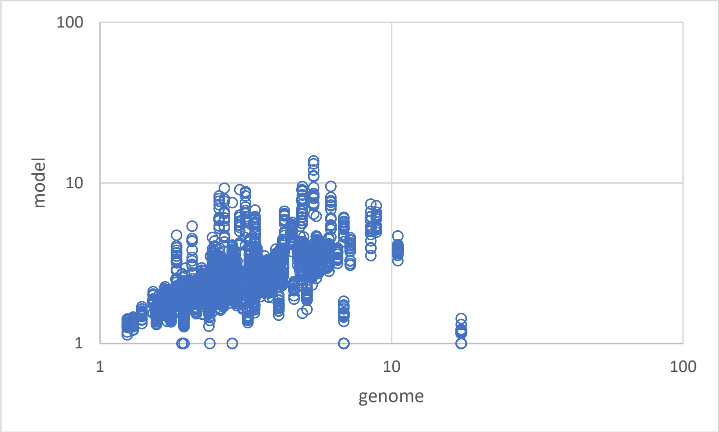


enrichment

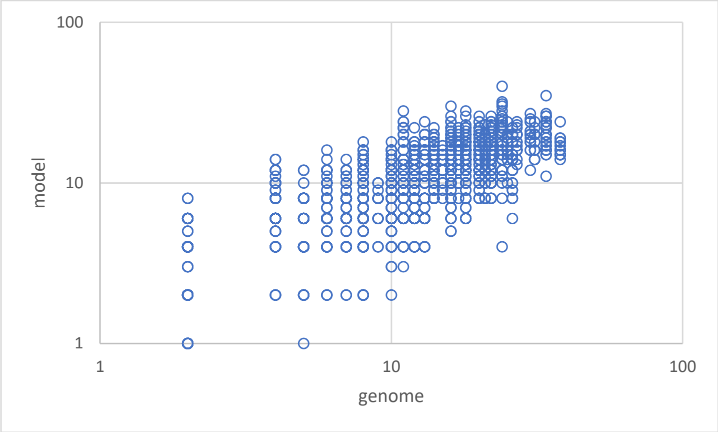


no. of segments

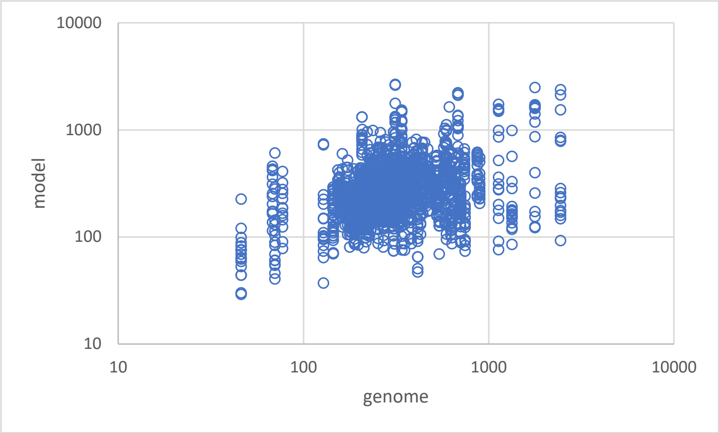


enriched segment length

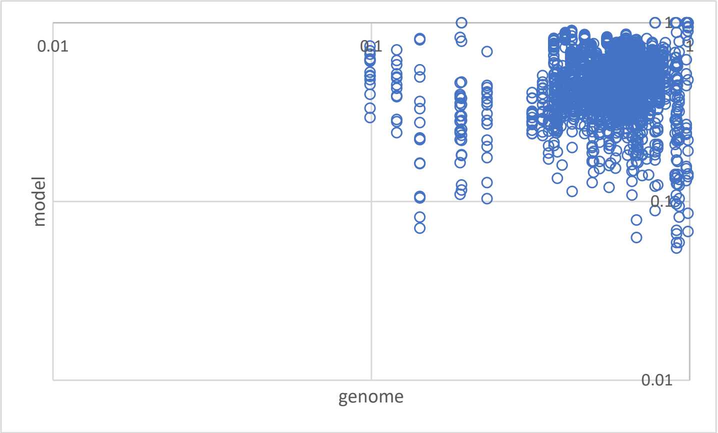


fraction of genome in enriched segments

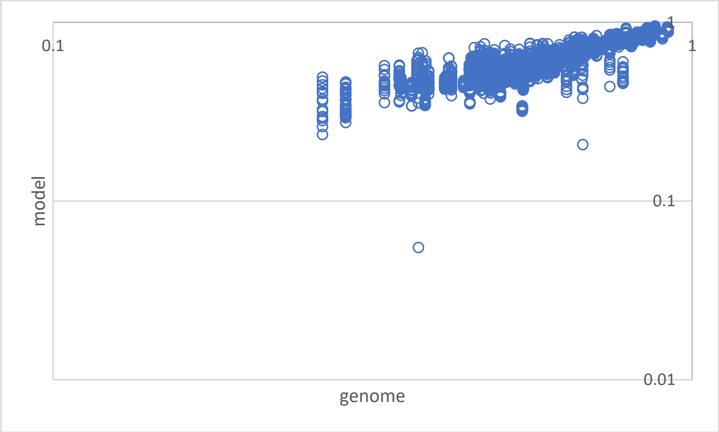


density of enriched segments

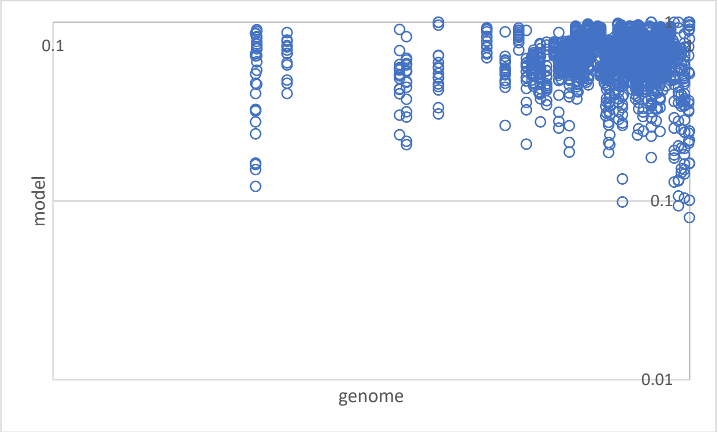


fraction of ancient genes in enriched segments
